# Evaluating hippocampal replay without a ground truth

**DOI:** 10.1101/2022.12.12.520040

**Authors:** M. Takigawa, M. Huelin Gorriz, M. Tirole, D. Bendor

## Abstract

During rest and sleep, memory traces replay in the brain. The dialogue between brain regions during replay is thought to stabilize labile memory traces for long-term storage. However, because replay is an internally-driven, spontaneous phenomenon, it does not have a ground truth - an external reference that can validate whether a memory has truly been replayed. Instead, replay detection is based on the similarity between the sequential neural activity comprising the replay event and the corresponding template of neural activity generated during active locomotion. If the statistical likelihood of observing such a match by chance is sufficiently low, the candidate replay event is inferred to be replaying that specific memory. However, without the ability to evaluate whether replay detection methods are successfully detecting true events and correctly rejecting non-events, the evaluation and comparison of different replay methods is challenging. To circumvent this problem, we present a new framework for evaluating replay, tested using hippocampal neural recordings from rats exploring two novel linear tracks. Using this two-track paradigm, our framework selects replay events based on their temporal fidelity (sequence-based detection), and applies a cross-validation using each event’s trajectory discriminability, where sequenceless decoding across both tracks is used to quantify whether the track replaying is also the most likely track being reactivated.

## Introduction

The hippocampus plays a central role in the encoding and consolidation of new memories **(Eichenbaum, 2000; Frankland and Bontempi, 2005; Klinzing et al., 2019; Lewis and Bendor, 2019; Squire, 1992)**. During locomotion in rodents, hippocampal place cells are active in specific regions of the animal’s environment (place fields), resulting in a sequential pattern of place cell firing when the animal runs along a spatial trajectory **(O’Keefe and Dostrovsky, 1971)**. During offline states, such as quiet restfulness and non-REM sleep, the sequential pattern of neural activity observed during behavior is spontaneously reactivated, a phenomenon referred to as “hippocampal replay” **(Lee and Wilson, 2002; Wilson and McNaughton, 1994)**. Offline replay of a neural memory trace is postulated to be a central mechanism by which memories recently encoded in the hippocampus can be consolidated by further stabilizing these memories in distributed cortico-hippocampal circuits for long-term storage **(Buzsaki, 1989; Klinzing et al., 2019).**

Evidence of replay was first discovered almost 30 years ago with the demonstration that during sleep, temporal correlations can be observed between the co-firing of place cell pairs with overlapping place fields **(Wilson and McNaughton, 1994)**. Since then, methods for large-scale chronic extracellular neural recordings have become more advanced, allowing the simultaneous recording of tens and even hundreds of neurons **(Ji and Wilson, 2007; Lee and Wilson, 2002; Pfeiffer and Foster, 2013; Widloski and Foster, 2022).** In line with these developments, the analysis methods for replay have also become more sophisticated - shifting from the pairwise analysis of place cells to detecting sequential patterns within neuronal ensembles, using either spiking activity directly **(Foster and Wilson, 2006)** or decoding this activity to extrapolate the virtual spatial trajectories replaying **(Davidson et al., 2009; Zhang et al., 1998).** Some of the most commonly used replay scoring metrics for quantifying the fidelity of a replay sequence include - 1) a *Spearman’s rank-order correlation* of spike times, which quantifies the ordinal relationship between the temporal order of place cell firing during behavior and a replay event’s spike train **(Foster and Wilson, 2006)**, but assumes that the place cell sequence is ordered accordingly to each place cell’s peak firing rate location alone, 2) a *Weighted correlation* of the decoded replay event, which quantifies a generalized linear correlation in time and position weighted by the decoded posterior probabilities without any assumption about the temporal rigidity of the replayed trajectory **(Grosmark and Buzsaki, 2016; Silva et al., 2015; Tirole & Huelin Gorriz, et al., 2022)**, and 3) a *Linear fitting* of the decoded replay event, which finds the linear path with the maximum summed decoded probability, assuming that the trajectory’s slope is constant **(Davidson et al., 2009; Gomperts et al., 2015; Ólafsdóttir et al., 2017).** However, because replay is generated by an internal and spontaneous state of the brain, there is no external reference to indicate whether a given replay event is truly a reinstatement of a memory trace. Without a ground truth, the detection of replay events must be inferred based on whether the statistical likelihood of observing a match between the sequential structure of the replayed event and the original behavioral template is sufficiently low, by chance. To quantify this, each event’s replay score is compared to a distribution of scores obtained using randomized data, permutated in either the spatial or temporal domain (i.e. a shuffled distribution), where statistically significant replay scores must be greater than a certain percentage of this shuffled distribution, typically 95% for a p-value < 0.05.

While recent replay studies have relied predominately on these three major methods of scoring replay (*rank-order, weighted correlation*, or *linear fit*), there are still many variations in how these scores can be calculated and how the subsequent statistical significance of these scores are measured. This can lead to issues in reproducibility and a greater difficulty in interpreting conflicting results between studies **(Tingley and Peyrache, 2020).** To overcome this problem, we need the ability to cross-validate replay events (i.e. are the real events being correctly detected, and non-events being correctly rejected), in spite of not having a ground truth **(Tingley and Peyrache, 2020; van der Meer et al., 2017; van der Meer et al., 2020).** Hypothetically, any method of cross-validation should at the very least be able to pass a basic test of distinguishing between replay events detected from real data and spurious replay events detected from randomized data.

Given that we are detecting a replay sequence using a replay score (and therefore p-value), we cannot use similar metrics for cross-validation due to a lack of independence. Here we solve this problem by cross-validating replay events using a sequenceless decoding approach, which has been used in several recent replay studies when the rat’s behavior involved running different trajectories, either on a T-maze with two arms or on two different linear tracks **(Carey et al., 2019; Tirole & Huelin Gorriz, et al., 2022)**. This approach quantifies how well the decoder can discriminate between two trajectories, with a greater difference in summed trajectory likelihoods indicating a higher discriminability. Based on replay data from rats running on two novel tracks, we validate this framework and demonstrate how measures of sequence fidelity combined with measures of trajectory discriminability provide the means to evaluate the performance of different replay detection strategies and methods.

## Results

Extracellular signals from the dorsal CA1 region of the hippocampus were recorded in rats running back and forth on two novel linear tracks (male Lister-hooded, n=5, 10 sessions, dataset from **Tirole & Huelin Gorriz, et al. (2022)**). In addition to running on two linear tracks (RUN), rats also had a rest/sleep session in a remote location both before (PRE) and afterwards (POST) (**Figure 1A**). To demonstrate how this framework quantifies replay detection performance in terms of dataset-specific detection rate, false-positive rate and trajectory discriminability, we started by detecting replay events using the weighted correlation of the decoded event, a common sequence-based replay detection approach **(Grosmark and Buzsaki, 2016)**.

**Figure 1.**
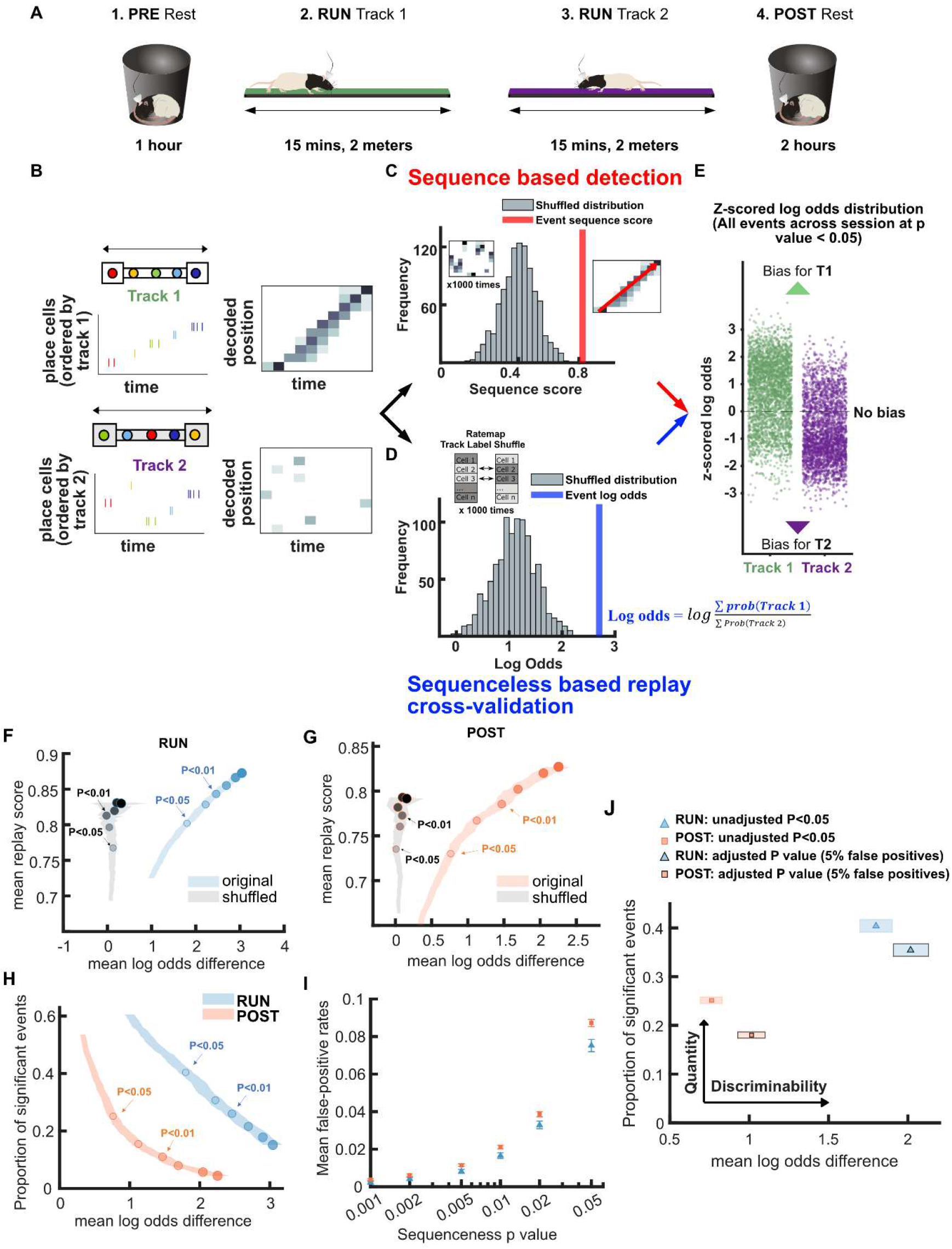
Demonstration of novel replay analysis framework for comparing sequence fidelity with trajectory discriminability. **(A)** Experimental design. For each recording session, the animal ran back and forth on two novel linear tracks (RUN) with resting sessions before (PRE) and afterwards (POST). **(B-E)** Schematic of sequence-based and sequenceless decoding framework. **(B)** Each candidate replay event spike train was fed into a naïve Bayesian decoder to calculate the decoded posterior probabilities across time and space. **(C)** Then, for the sequence-based analysis, the sequence score for each candidate event was determined from the weighted correlation of the posterior probability matrix. Significance was determined by comparing the replay score relative to a shuffled distribution (p-value < 0.05). **(D)** For sequenceless decoding, a Bayesian decoding similar to sequence-based approach was used, with the exception that only place cells with stable place fields on both tracks were used as template to avoid any track discrimination bias. Then, the logarithmic ratio of the summed posterior probabilities within each replay event for each track is calculated (log odds). The event log odds were z-scored relative to a shuffled distribution where each place cell’s track 1 and 2 place fields were randomly shuffled between tracks. **(E)** The difference between track 1 and track 2 replay events’ log odds can be used as a metric to cross-validate the performance of sequence-based replay detection. **(F,G)** The relationship between the mean log odds difference and mean replay score for the significant events detected at different p-value thresholds (0.2 to 0.001) using weighted correlation replay scoring with two different shuffling procedures (place field circular shuffle and time bin permutation shuffle). The shaded region indicates the 95% bootstrap confidence interval for the mean log odds difference. The six dots with increasing color intensity for each distribution represent the data at a p-value threshold of 0.05, 0.02, 0.01, 0.005, 0.002 and 0.001. **(F)** Significant events detected during RUN using original candidate events (blue) and cell-id shuffled spurious events (grey). **(G)** Significant events detected during POST using original candidate events (orange) and cell-id shuffled spurious events (grey). **(H)** The relationship between the mean log odds difference and the proportion of significant events detected at different p-value thresholds (0.2 to 0.001) during RUN (blue) and POST (orange). The shaded region indicates the 95% bootstrap confidence interval for mean log odds difference. The six dots with increasing color intensity for each distribution represent the data at a p-value threshold of 0.05, 0.02, 0.01, 0.005, 0.002 and 0.001. **(I)** The mean proportion of significant cell-id shuffled events (mean false-positive rate) at different p-value thresholds (0.2 to 0.001). The error bar indicates the 95% bootstrap confidence interval for mean false-positive rates. **(J)** The replay detection performance at the original p < 0.05 and an adjusted p-value when the mean false-positive rate was 5%. The shaded box indicates a 95% bootstrap confidence interval for both proportion of significant events detected and mean log odds difference. The box with a light outline represents the values at p < 0.05 and the box with black outline represents the values at the adjusted p-value. The 95% confidence interval for the proportion of significant events, mean log odds difference, mean false-positive rates and the adjusted p-value for replay events detected during RUN and POST are available in Figure 1—source data 1.

Candidate replay events were population burst events (peak z-scored multi-unit activity >3) that occurred during periods of inactivity, when the animal’s velocity is less than 5 cm/s (see Methods). Across 10 sessions, we detected 8485, 4643 and 14326 candidate events in PRE, RUN and POST session, respectively. Detected replay events were required to also have a ripple power z-score greater than 3 and a significant replay score - that is, higher than 95% of the scores obtained from each of two shuffle distributions, namely a place field circular shuffle and a time bin permutation shuffle (see Methods) **(Grosmark and Buzsaki, 2016).** The replay score was obtained by performing a weighted correlation on the posterior probabilities (across place and time) obtained using a naïve Bayesian decoder **(Figure 1B-C)**. Because the dataset includes more than one track, the posterior probabilities across the two available tracks were normalized at each time bin such that their combined sum was one (**Figure 1B) (Bendor and Wilson, 2012; Carey et al., 2019).**

### Demonstrating novel framework for cross-validation of replay detection performance

We next created a framework for evaluating and comparing replay detection methods, using a comparison between a replay event’s sequence fidelity and its trajectory discriminability. For each replay event detected, we used a sequenceless, log odds metric *based on only place cells with place fields on both tracks*, to avoid any potential bias in track discrimination (**Figure 1D-E** see Methods) **(Carey et al., 2019; Tirole & Huelin Gorriz, et al., 2022)**. Sequenceless decoding involved three steps: 1) computing the summed posteriors (across time bins) within the replay event for each track, 2) calculating the ratio between the log of the summed posteriors for each track, and 3) taking the z-score of this value, based on a distribution of log odds computed by a track ID shuffle **(Carey et al., 2019; Tirole & Huelin Gorriz, et al., 2022)**. For each place cell in the track ID shuffle, the corresponding track 1 and 2 place fields were randomly assigned to the correct track or swapped. As a result, a more positive z-scored log odds would indicate a greater likelihood of track 1 reactivation whereas a more negative value would indicate a greater likelihood of track 2 reactivation (**Figure 1E**). To use sequenceless decoding to cross-validate replay events, we computed the difference in mean log odds between track 1 and track 2 replay events, which were originally detected using standard sequence-based replay detection methods. For this measurement, a more positive mean log odds difference would indicate a higher trajectory discriminability in the replay content, and a higher confidence in replay quality. In contrast, a mean log odds difference of 0 would suggest that the quality of replay events is comparable to chance level trajectory discriminability, and that the detected event population most likely consists of low fidelity events, indistinguishable from false-positives.

As mentioned earlier, any method of cross-validation should at least be able to pass a basic test of distinguishing between replay events detected from real data and spurious replay events detected from randomized data. We first compared the mean log-odds difference and mean weighted correlation score for replay events detected using our dataset and spurious replay events detected after the same dataset was randomized using a cell-id shuffle. Note that all sequence-based replay detection methods in this study did not use a cell-id shuffle, and as such this shuffling approach was sufficiently independent for generating random sequences as a negative control. When p-value threshold was varied from 0.2 to 0.001, we observed a close relationship between mean weighted correlation score and p-value threshold for both the original and randomized dataset (**Figure 1F,G**). However, there was a clear dissociation between p-value threshold and mean log odds difference when using shuffled dataset, for both RUN and POST replay events. This demonstrated that while weighted correlation score improved with p-value thresholds even for shuffled dataset, the mean log odds difference (trajectory discriminability) was able to differentiate real from spurious replay events, acting independently from our selection criteria of replay events based on sequence fidelity.

While a p-value of 0.05 is usually the chosen threshold for whether a statistical test rejects the null hypothesis, we predicted that as the p-value criteria became stricter, the mean log odds difference should increase due to a lower rate of false-positive events, albeit at the cost of also fewer detected replay events overall. To study the trade-off between the trajectory discriminability and detection rate of detected replay events, we first analyzed our POST and RUN replay events, comparing the number of significant events detected on both tracks to the mean log odds difference, as the p-value threshold was adjusted between 0.2 to 0.001 (**Figure 1H**). For both POST and RUN, the mean log odds difference increased (higher trajectory discriminability) as the p-value threshold decreased, which also corresponded to a decrease in the number of detected events. In addition, we observed RUN replay events to be comparatively more prevalent and yielding a higher trajectory discriminability than POST replay events, in line with previous reports **(Karlsson and Frank, 2009; Tirole & Huelin Gorriz, et al., 2022)**. This might be partly due to the presence of immediate sensory or other external inputs helping to direct hippocampal place cell ensembles towards representing the local current environment during awake replay.

To determine if the decrease in detection rate was associated with decrease in the false-positive rate for this detection method, we calculated the false-positive rate (mean fraction of spurious replay events detected across both tracks) using the full range of p-value thresholds tested (from 0.2 to 0.01). We observed that as p-value threshold was gradually reduced, so was the false-positive rate (**Fig. 1D**). We also found that the empirically-measured false-positive rate was higher than the p-value: using a p-value of 0.05, 7.5% (lower CI = 7.2% and higher CI = 7.8%) of RUN replay events were false-positives, while 8.7% (lower CI = 8.5% and higher CI = 8.9%) of POST replay events were false-positives. Using this information, we could quantify the proportion of significant events for both tracks, and the corresponding mean log odds difference at the original p-value of 0.05 and an adjusted p-value (RUN: p=0.032, POST: p=0.028) that corrected the mean false-positive to approximately 5% (**Fig 1E**). Such adjustment of p-value based on equivalent false-positive rate would provide an opportunity to compare different replay detection strategies and methods in a more equitable manner.

We next used our framework to analyze and compare different strategies for replay detection. We did not attempt to precisely replicate any specific published replay detection method, as our goal was to see how general methodological differences can impact trajectory discriminability and quantity, and whether there is a preferred general approach for replay analysis. While it is important to note that there are many subtler, data-specific details to potentially consider (e.g. number of cells, smoothing of data, bin size, etc.) that can impact both the false-positive rate and quality of detected replay events, our aim was to provide an overarching framework to help direct us to a more optimal replay detection method.

### Ripple power is an important criterion for replay event selection during POST

During both quiet wakefulness and non-REM sleep, replay events preferentially occur during sharp-wave ripple events, transient high frequency oscillations (150-250 Hz) in the local field potential (LFP) recorded near the cell layer of CA1 **(Buzsaki, 2015; Foster and Wilson, 2006; Lee and Wilson, 2002).** As such, a minimum ripple power has been used as a criterion for detecting candidate replay events prior to sequence detection, however the threshold has varied substantially across previous studies ranging from 2 SD (Standard Deviations) above baseline **(Diba and Buzsaki, 2007; Gillespie et al., 2021)** to 7 SD above baseline **(Csicsvari et al., 2007; Ji and Wilson, 2007).** However, because many replay events consist of the spontaneous reactivation of a large proportion of place cells within a short time window (more than typically occurs during active behavior), a substantial increase in multiunit activity during immobility also provides a reasonable criterion for detecting candidate replay events **(Davidson et al., 2009).** Because using only multiunit activity as a threshold typically leads to more candidate replay events, and is believed to be more reliable for detecting both the start and end times of the replay event, many recent studies have opted to no longer use minimum ripple power as a criterion for candidate replay events **(Gomperts et al., 2015; Ólafsdóttir et al., 2018; Olafsdottir et al., 2016; Silva et al., 2015).**

We next applied our replay analysis framework to test how a stricter criterion for ripple power affects trajectory discriminability. Candidate events were selected based on both elevated MUA (z-score>3) and ripple power limited to a specific range, measured in SD above baseline (i.e. a z-score of 0-3, 3-5, 5-10, or >10). For matched p-values, a stricter ripple threshold resulted in both a higher proportion of significant events and a higher mean log odds difference(**Figure 2A,B**). However, one major difference between RUN and POST replay events is that even at the lowest ripple power threshold, RUN replay events had a non-zero mean log odds difference, which increased with stricter p-values. In contrast for POST replay events with a ripple power less than a z-score of 5, the mean log odds difference was near zero for all p-value thresholds tested. Furthermore, while the proportion of detected POST events with a ripple power less than a z-score of 5 was approximately 20% at p-value of 0.05, the proportion of detected POST events dropped to near chance levels of 10% when p-value was adjusted to match the mean false-positive rate of 5% (where 10% of spurious events were detected as significant events across both tracks) (**Figure 2C,D**, see method). For ripple thresholds above 5, the log odds difference increased as the proportion of events and p-value decreased **(Figure 2A,B)**. The increase in trajectory discriminability observed with higher ripple power was not associated with a change in the number of active place cells or total place cell spiking activity **(Figure 2 SUPP1).** These results emphasize that a ripple power threshold is not necessary for RUN replay events but may still be beneficial, as long as it does not excessively eliminate too many good replay events with low ripple power. In other words, it is possible that a stricter p-value with no ripple threshold can still detect more replay events than using a less strict p-value combined with a strict ripple power threshold. However, for POST replay events, a threshold at least in the range of a z-score of 3-5 is necessary, to reduce inclusion of false-positives within the pool of detected replay events. For the remainder of this study, we incorporated a strict, albeit less conservative ripple power threshold (z-score>3) for both RUN and POST, given the trade-off between an improved trajectory discriminability and a decrease in the number of detected events. This also allowed a more direct comparison of awake and rest replay using identical criteria while still being consistent with previous replay studies **(Karlsson and Frank, 2009; Pfeiffer and Foster, 2013; Todorova and Zugaro, 2019)**

**Figure 2.**
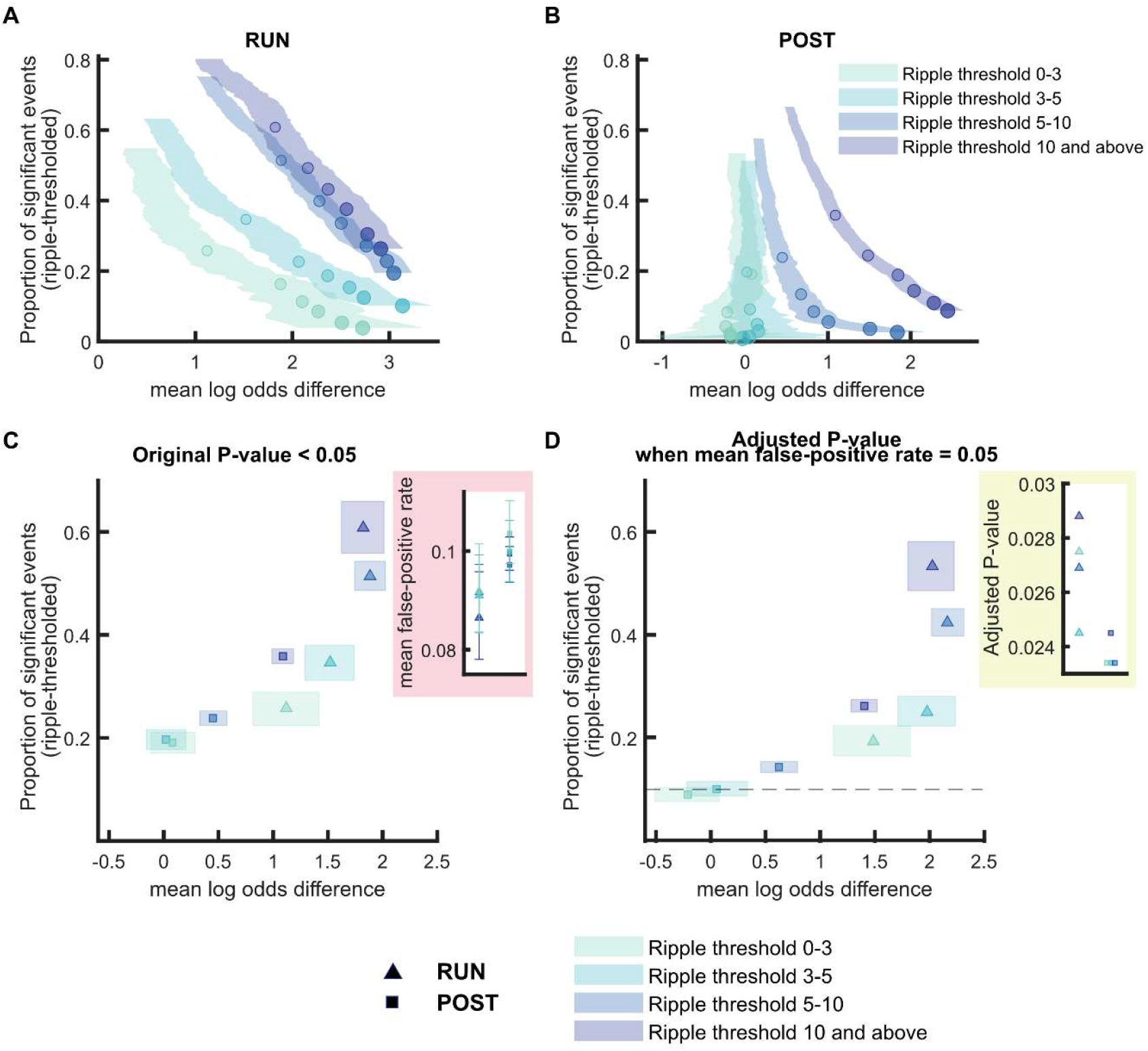
Replay detection performance improves with ripple power. **(A,B)** The proportion of significant events and mean log odds difference at different p-value thresholds (0.2 to 0.001) as ripple power increases (0-3, 3-5, 5-10, 10 and above). The shaded region indicates the 95% bootstrapped confidence interval for mean log odds difference. The six dots with increasing color intensity for each distribution represent the data at a p-value threshold 0.05, 0.02, 0.01, 0.005, 0.002 and 0.001. **(A)** Replay events detected during RUN. **(B)** Replay events detected during POST. **(C,D)** The proportion of significant events and mean log odds differences at **(C)** an unadjusted p-value < 0.05 and **(D)** an adjusted p-value with a mean false-positive rate = 5%. The shaded box indicates a 95% bootstrap confidence interval for both the proportion of significant events detected and mean log odds difference. The triangle symbol is used to represent replay events during RUN and the square symbol is used to represent replay events during POST. The dashed line represents the approximate chance level at mean false-positive rate of 5%. The 95% confidence interval for the proportion of significant events, mean log odds difference, mean false-positive rates and the adjusted p-value for replay events with different ripple power range are available in Figure 2—source data 1.

### Replay detection is sensitive to the shuffle method used

Statistical significance of each replay event is calculated by comparing the replay score to one or more Monte-Carlo shuffle distributions, with each shuffle designed to randomize a specific aspect of sequential place cell firing while attempting to keep other factors intact. Some shuffles are applied to randomize the place fields or the spike train of the original data prior to Bayesian decoding, while other shuffles are applied to the spatial or temporal dimension of the decoded posterior probabilities. Because place cells often fire bursts of spikes, this creates non-independent samples, which *violate the assumption of having independent samples for a statistical test*. Thus, it is important for a shuffle method to preserve such aspects of the data that are not independent, to avoid adding type 1 errors in the statistical analysis, leading to a false-positive rate that exceeds the p-value. Given this, we next examined how different shuffling procedures impact replay detection performance. Using the weighted correlation scoring method, we examined four types of shuffling procedures to see if they differed in how they detected replay during RUN and POST epochs. These consisted of two pre-decoding shuffles (spike train circular shuffle, place field circular shuffle) and two post-decoding shuffles (place bin circular shuffle, time bin permutation shuffle) **(Figure 3A-D, Figure 3 SUPP)**.

**Figure 3.**
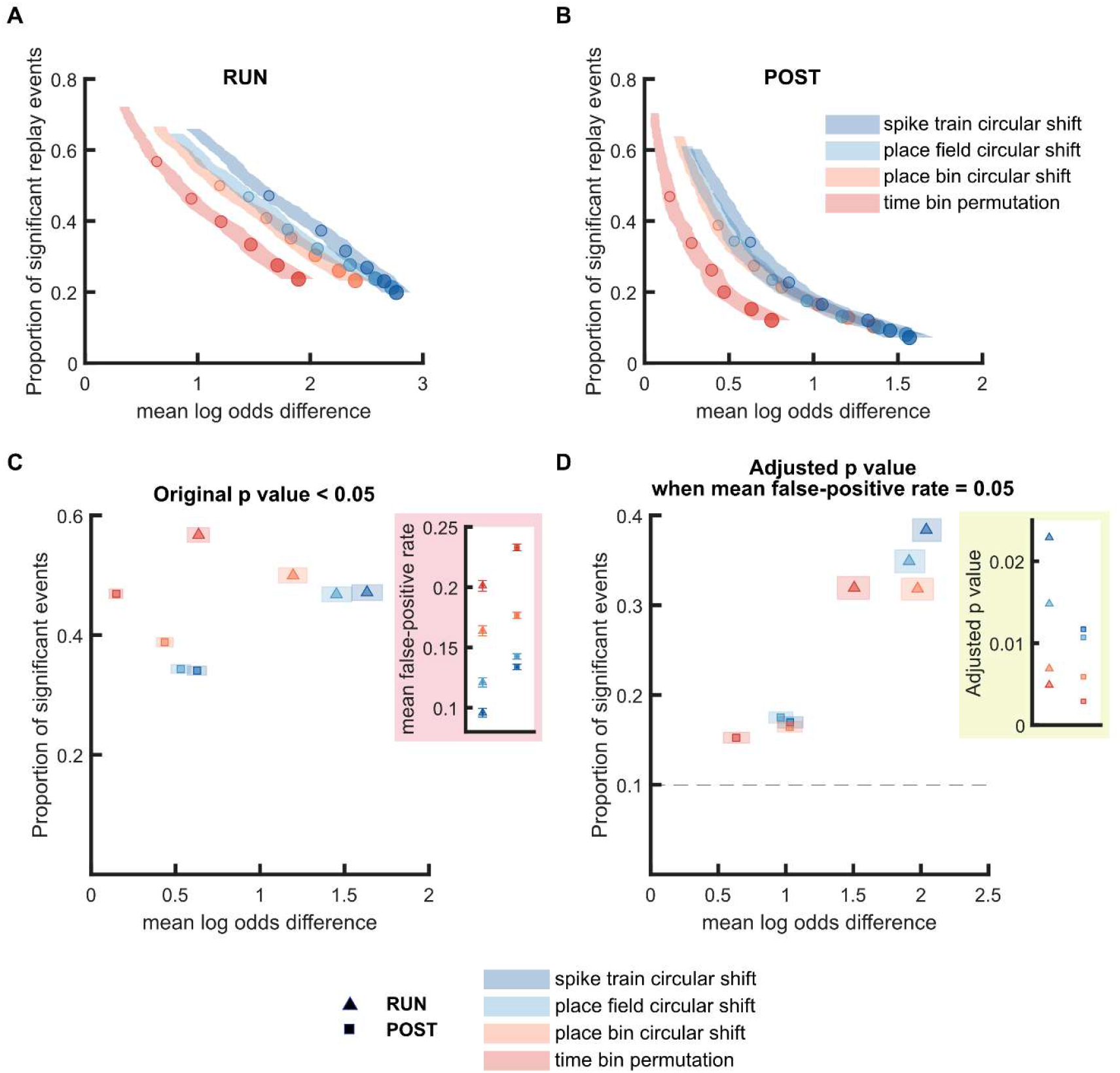
Replay detection performance was sensitive to the shuffling method applied. **(A,B)** The proportion of significant events and mean log odds difference at different p-value thresholds (0.2 to 0.001) when using four different shuffling methods: **(1)** spike train circular shuffle (dark blue), **(2)** place field circular shuffle (light blue), **(3)** place bin circular shuffle (orange) and **(4)** time bin permutation shuffle (red). The shaded region indicates the 95% bootstrapped confidence interval for mean log odds difference. The six dots with increasing color intensity for each distribution represent the data at p-value threshold 0.05, 0.02, 0.01, 0.005, 0.002 and 0.001. **(A)** Replay events detected during RUN. **(B)** Replay events detected during POST. **(C,D)** The proportion of significant events and mean log odds difference at **(C)** an unadjusted p-value < 0.05 and **(D)** an adjusted p-value with a 5% false-positive rate. The shaded box indicates the 95% bootstrap confidence interval for both the proportion of significant events detected and mean log odds difference. The triangle symbol is used to represent replay events during RUN and the square symbol is used to represent replay events during POST. The dashed line represents the approximate chance level at mean false-positive rate of 5%. The 95% confidence interval for the proportion of significant events, mean log odds difference, mean false-positive rates and the adjusted p-value for replay events detected using different shuffling methods are available in Figure 3—source data 1.

For both RUN and POST, we observed that pre-decoding shuffles performed better than post-decoding shuffles, with the time bin permutation shuffle showing a significantly lower log odd difference compared to the other three shuffles (**Figure 3A,B**). Although the post-decoding time bin permutation shuffle seemingly led to the highest detection rate at a p-value of 0.05 (**Figure 3C**), we observed that its mean false-positive rate at a p-value of 0.05 was also substantially higher than other shuffles. When we adjusted the p-value to match the mean false-positive rate of 5%, the post-decoding time bin permutation shuffle had the poorest performance (the lowest proportion of significant events and the lowest mean log odds difference) across the four types of shuffles for both RUN and POST epochs (**Figure 3D**). For both the RUN and POST epochs, the remaining three shuffles had a similar mean log odds difference when using an adjusted p-value. However, while the proportion of significant events were similar across the three shuffles for POST, the two pre-decoding shuffles had a higher proportion of significant events for RUN replay events. These results suggested that the shuffling procedures that directly manipulated the place field data or spike train data before decoding may be more efficacious than these procedures that randomized the posterior probability distributions after decoding.

Now that we have examined the merits of individual shuffles, we next evaluated how replay detection performance might be further improved.

### Replay detection can be improved by adding stricter detection criteria

Rather than lowering the false-positive rate by using a stricter p-value, another approach is to apply additional criteria to the selection of a significant replay event. Commonly this is accomplished by using multiple types of shuffling procedures, with the requirement that each shuffling method must independently pass the same p-value criterion for the replay event to be classified as statistically significant. An alternative approach to using multiple shuffling procedures is to impose a criteria of jump distance - the number of spatial bins the decoded estimated position is allowed to jump across neighboring time bins during a spatial trajectory, with candidate replay events rejected if they fail this criterion **(Silva et al., 2015).** One justification for using this metric is that a lower jump distance is correlated with a higher weighted correlation score, however not all events with higher scores are necessarily higher quality events.

Based on this, we compared the performance of four replay detection methods varying in their shuffling procedures-1) a single place field circular shuffle, 2) a place field circular shuffle with a jump distance threshold (40% of track) **(Silva et al., 2015)**, 3) two shuffles (place field circular shuffle and time bin permutation shuffle) **(Grosmark and Buzsaki, 2016)**, and 4) three shuffles (place field circular shuffle, spike train circular shuffle and place bin circular shuffle) **(Tirole, Huelin-Gorriz, et al. 2022)**. For both RUN and POST epochs, increasing the number of shuffles increased the mean log odd difference and decreased the proportion of significant events and false-positive rate, for similar p-values (**Figure 4A,B and Figure 4A,B SUPP1**). Adding a jump distance threshold as an additional criterion also increased the mean log odd difference, although this was accompanied by a marked reduction in the proportion of significant replay events (**Figure 4A,B**). When using an unadjusted p-value, two methods (1 shuffle with a jump distance threshold and 3 shuffles) resulted in the highest mean log odd difference (**Figure 4C**). However, if an adjusted p-value was used (with the false-positive rate fixed at 0.05), only the use of 3 shuffles achieved both the highest proportion of events and largest mean log odds difference, for both RUN and POST epochs (**Figure 4D**). In contrast to this, a single shuffle with an added jump distance criterion performed the poorest (less events and lower trajectory discriminability) out of the four methods when using an adjusted p-value. This suggests that it is more beneficial to use a stricter p-value than to impose a jump distance, an observation that extended to a range of different jump distances spanning 20% to 60% of the track (**Figure 4 SUPP2**).

**Figure 4.**
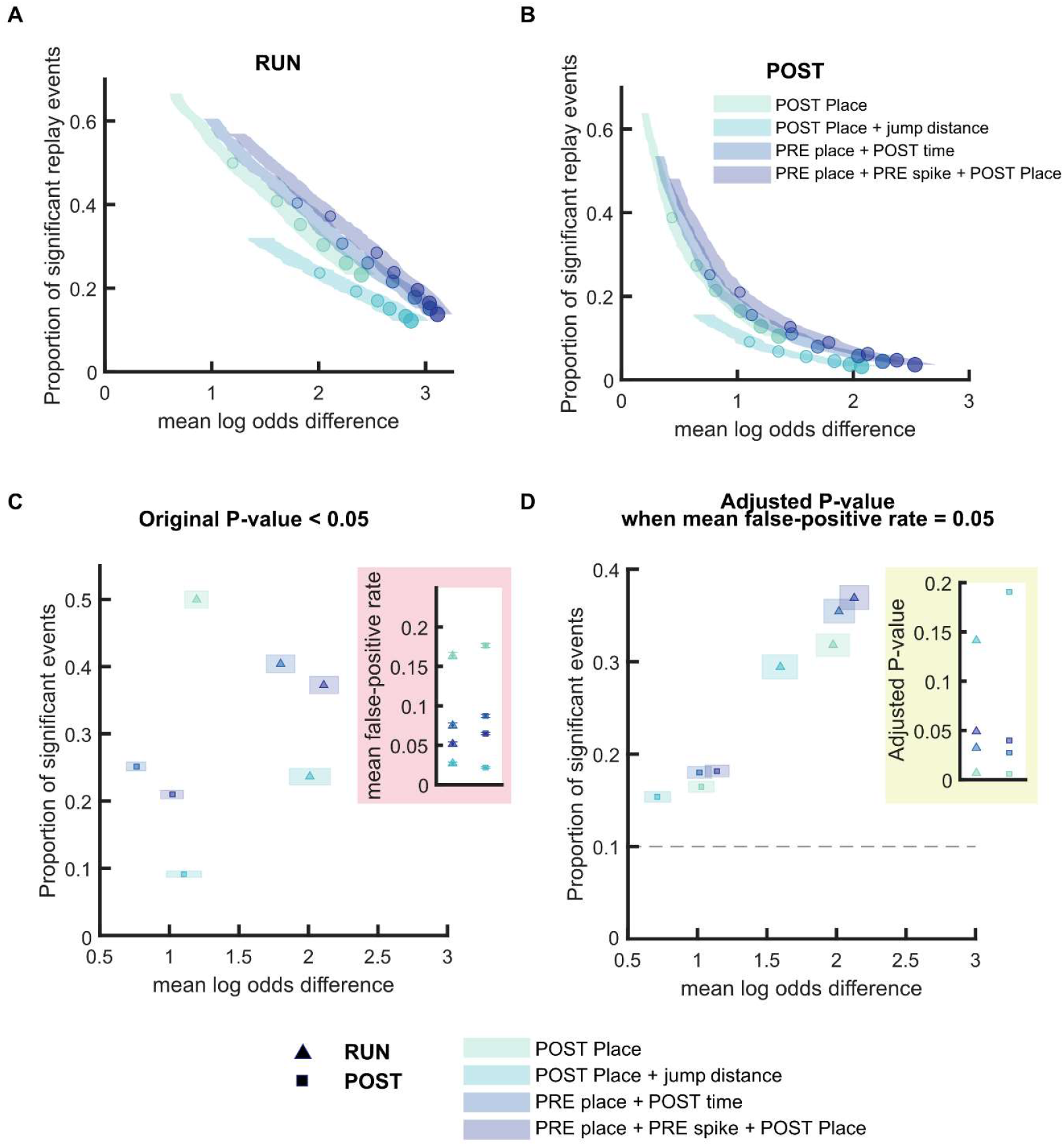
Replay detection performance can be improved by adding stricter detection criteria. **(A,B)** The proportion of significant events and mean log odds difference at different p-value thresholds (0.2 to 0.001) with four different detection criteria: **(1)** Only a post-decoding place bin circular shuffle **(2)** a post-decoding place bin circular shuffle with jump distance threshold at normalized track length 0.4 **(3)** a pre-decoding place field circular shuffle and a post-decoding time bin permutation shuffle, **(4)** a pre-decoding place field circular shuffle, a pre-decoding spike train circular shuffle and a postdecoding place bin circular shuffle. The shaded region indicates a 95% bootstrap confidence interval for mean log odds difference. The six dots with increasing color intensity for each distribution represent the data at p-value threshold 0.05, 0.02, 0.01, 0.005, 0.002 and 0.001. **(A)** Replay events detected during RUN. **(B)** Replay events detected during POST. **(C,D)** The proportion of significant events and mean log odds difference at **(C)** an unadjusted p-value < 0.05 and **(D)** an adjusted p-value with a 5% false-positive rate. The shaded box indicates a 95% bootstrap confidence interval for both the proportion of significant events detected and mean log odds difference. The triangle symbol represents replay events during RUN and the square symbol represents replay events during POST. The dashed line represents the approximate chance level at mean false-positive rate of 5%. The 95% confidence interval for the proportion of significant events, mean log odds difference, mean false-positive rates and the adjusted p-value for replay events detected using different detection criteria are available in Figure 4—source data 1.

### Rank-order based replay detection is sensitive to spike selection

We next extended our framework to investigate other replay scoring methods – first focusing on the use of rank-order correlation. Instead of scoring the decoded trajectories, this approach directly quantifies the ordinal relationship within the spiking patterns **(Foster and Wilson, 2006).** In this method, the sequential order of place fields is ranked along the track, and compared with the corresponding rank of the spike times within the replay event. Rank order correlation has been previously applied in two ways-1) to all spikes within the replay event **(Dragoi and Tonegawa, 2011; Foster and Wilson, 2006)** or 2) only to one spike emitted by each place cell during a candidate replay event (e.g. median or first spike) **(Tirole, Huelin-Gorriz, et al. 2022).** However, as mentioned previously, place cells often fire bursts of spikes which in effect creates non-independent samples. Therefore, in theory, the inclusion of all spikes for the Spearman’s rank-order based analysis would violate the statistical assumption of independent samples and lead to more type I errors (and a higher false-positive rate). We compared the rank-order based analysis using either all spikes or median spike in our framework, and found that the false-positive rate when using independent events (median spike time) remained around 5% (lower CI = 5.2% and upper CI = 5.8%), but increased to around 17% (lower CI = 16.3 and upper CI = 17.2%), when using all spikes (**Figure 5 and Figure 5 SUPP**). However, when using the adjusted p-value (falsepositive rate fixed at 5%), the trade-off between the proportion of significant events and their mean log odds difference was similar between the two methods (**Figure 5D and Figure 5 SUPP**).

**Figure 5.**
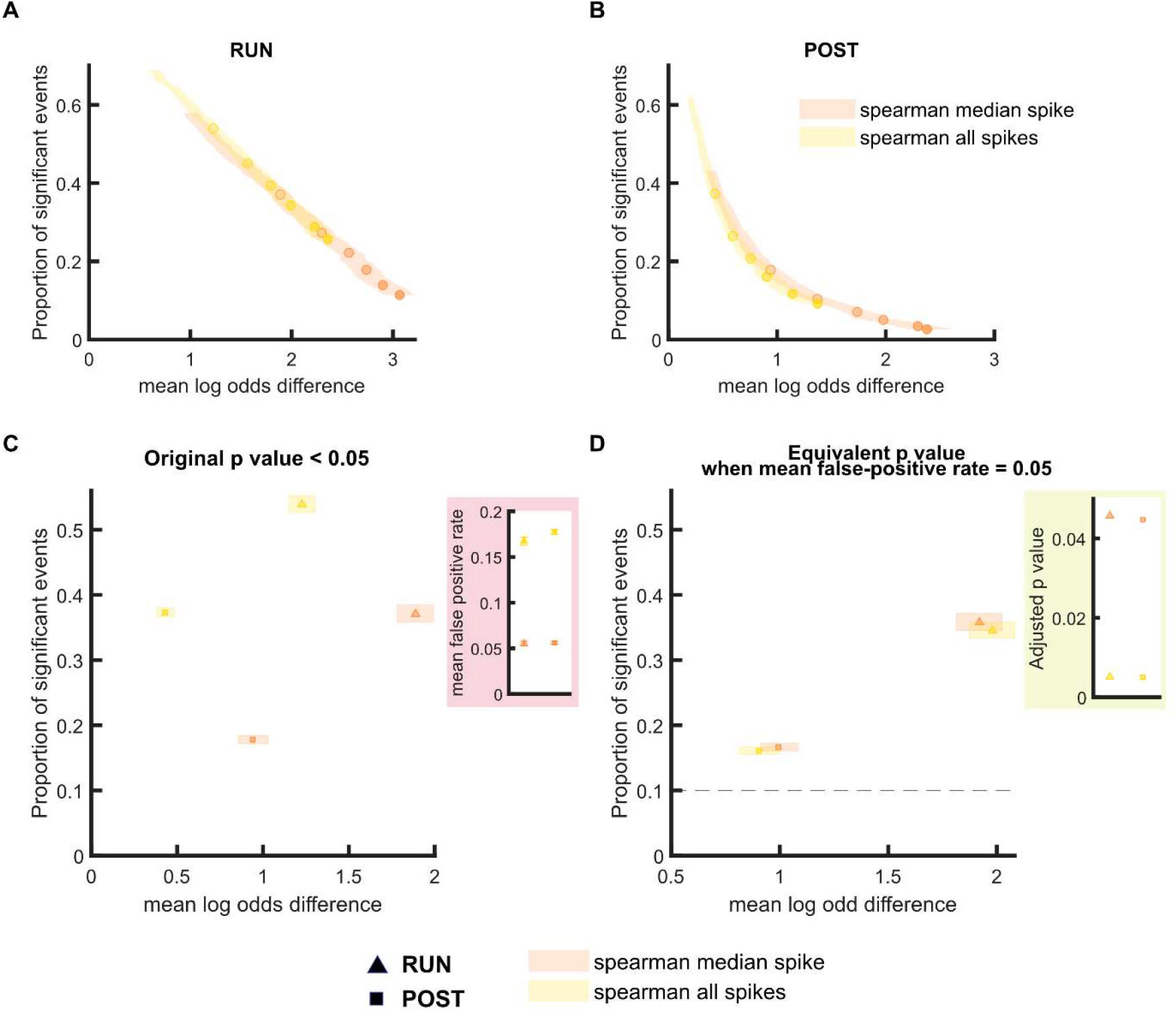
The performance of rank-order based replay detection method depends on the selection of spikes for analysis. **(A,B)** The proportion of significant events and mean log odds difference at different p-value thresholds (0.2 to 0.001) when **(1)** all spikes or **(2)** only the median spike fired by each place cell was included for rank-order based replay analysis. The shaded region indicated 95% bootstrap confidence interval for mean log odds difference. The six dots with increasing color intensity for each distribution represent the data at p-value threshold 0.05, 0.02, 0.01, 0.005, 0.002 and 0.001. **(A)** Replay events detected during RUN. **(B)** Replay events detected during POST. **(C,D)** The proportion of significant events and mean log odds difference at **(C)** an unadjusted p-value < 0.05 and **(D)** an adjusted p-value with a 5% false-positive rate. The shaded box indicates a 95% bootstrap confidence interval for both the proportion of significant events and mean log odds difference. The triangle symbol is used to represent replay events during RUN and the square symbol was is to represent replay events during POST. The dashed line represents the approximate chance level at mean false-positive rate of 5%. The 95% confidence interval for the proportion of significant events, mean log odds difference, mean falsepositive rates and the adjusted p-value for rank-order based methods using all spikes or only median spike within the replay event are available in Figure 5—source data 1.

### Replay events have a similar quality across different detection methods, when using an adjusted p-value, but vary in their proportion

Finally, we expanded our analysis to directly compare every replay scoring and shuffling method already implemented in this report, with the addition one more commonly used method - a linear fit based score of decoded trajectories with either one and two types of shuffles **(Davidson et al., 2009; Muessig et al., 2019; Ólafsdóttir et al., 2017)**. In the linear fit approach, the decoded posterior probability matrix, representing the probability of the virtual trajectory at each position on the track and at each time point, is compared to the best linear trajectory; the corresponding replay score is the sum of all posterior probabilities within a certain distance of this line, after maximizing this score across all possible lines that can pass through this matrix. While the linear fit approach is more stringent than the weighted correlation, it may also be less sensitive to non-linear replay trajectories (e.g. sigmoidal). When using a cell-id shuffle to examine the chance-level detection for each method, we observed no bias in the mean log odds difference for every method, except a linear fit approach with two shuffles (**Figure 6 SUPP1**). To avoid any possible intrinsic bias in our calculation of the mean log odds, we calculated the *adjusted mean log odds difference*, by subtracting any potential bias in log odds difference measured using spurious replay events (**Figure 6**). Similar results were observed using an alternative within-track normalization approach in Bayesian decoding prior to replay detection (**Figure 6 SUPP2**). Finally, to provide a more comprehensive comparison across methods, we also included replay events detected in the PRE epochs, a phenomenon referred to as de-novo preplay where statistically significant sequences are detected prior to any knowledge or experience running on the track **(Dragoi and Tonegawa, 2011)**.

Using our framework to compare replay detection methodologies, which incorporated **1)** the corrected mean log odds difference, **2)** the mean false-positive rate based on replay detection using cell-id shuffled data, and **3)** the proportion of significant events on both tracks, we observed four main findings regarding replay during three different behavioral epochs (**Figure 6 and Figure 6 SUPP3-5**). First, the proportion of significant events varied substantially across methods when using the unadjusted p-value, however this was consistent with the false-positive rate, which ranged from 2% to 18%. Second, using the adjusted p-value, all methods had a proportion of significant preplay events close to 10%, which was similar to the proportion of false-positive events expected by chance for two tracks. Furthermore, the shuffle-subtracted mean log odds difference for preplay was approximately zero for all methods, suggesting that regardless of the method used, if used correctly, preplay is indistinguishable from randomly generated spurious replay events. Third, using the adjusted p-value for POST replay events, all methods produced a corrected mean log odds difference near 1, however linear and weighted correlation methods with 2 or more shuffles detected a higher proportion of significant events. Finally, using the adjusted p-value for RUN replay events, the majority of methods produced a corrected mean log odds difference near 2. The best two methods were a weighted correlation with 3 shuffles and linear fit with 2 shuffles (the latter detecting more events but having a lower trajectory discriminability). Ultimately, minor variations to linear fitting or weighted correlation may improve such methods further, however both of these methods, as long as a sufficient number of shuffles are used (preferentially pre-decoding shuffles), provide the best method of replay detection across the different methods tested with our framework. It is likely that replay detection methods will be further improved in the coming years, however quantifying how much better a new method is will now be possible using the framework we outline here.

**Figure 6.**
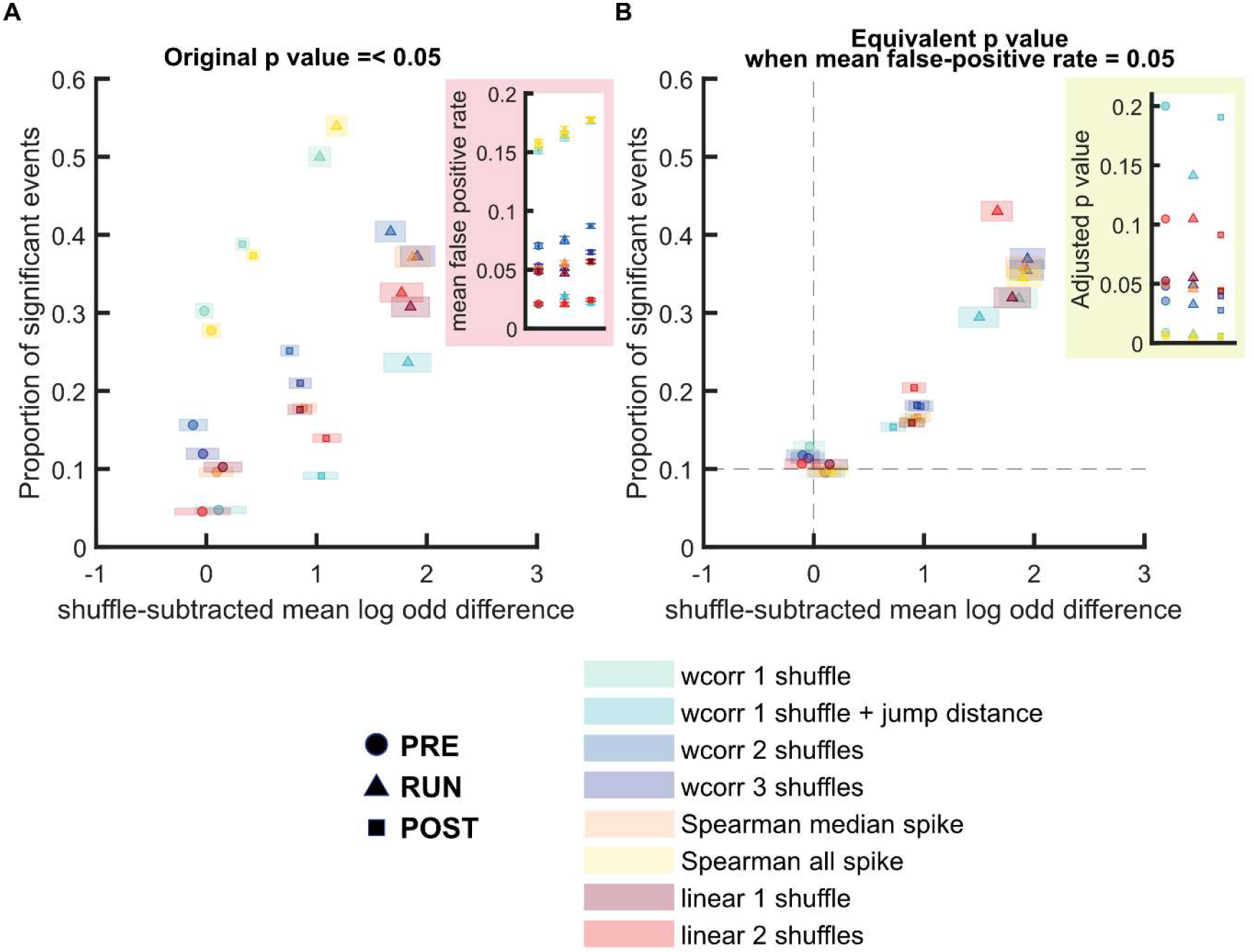
Comparison of different replay detection methods for replay events during PRE, RUN and POST. **(A,B)** The proportion of significant events and adjusted mean log odds difference at different p-value thresholds (0.2 to 0.001) using a range of different methods: **(1)** a weighted correlation replay scoring with place bin circular shuffle **(2)** a weighted correlation replay scoring with place bin circular shuffle and jump distance threshold at normalized track length 0.4 **(3)** a weighted correlation replay scoring with place field circular shuffle and time bin permutation shuffle, **(4)** a weighted correlation replay scoring with place field circular shuffle, spike train circular shuffle and time bin permutation shuffle. **(5)** a rank-order correlation using only the median spike time of each place cell **(6)** a rank-order correlation using all spikes within the event for all place cells **(7)** A linear fit replay scoring with place bin circular shuffle **(8)** a linear fit replay scoring with a place field circular shuffle and time bin permutation shuffle. The shaded region indicates a 95% bootstrap confidence interval. The six dots with increasing color intensity for each distribution represent the data at p-value threshold 0.05, 0.02, 0.01, 0.005, 0.002 and 0.001. **(A)** Replay events detected during RUN. **(B)** Replay events detected during POST. **(C,D)** The proportion of significant events and mean log odds difference at **(C)** an unadjusted p-value < 0.05 and **(D)** an adjusted p-value with a 5% false-positive rate. The shaded box indicates a 95% bootstrap confidence interval. The circle symbol is used to represent replay events during PRE. The triangle symbol is used to represent replay events during RUN and the square symbol is used to represent replay events during POST. The dashed line represented the approximate chance level detection rate and trajectory discriminability at mean false-positive rate of 5%. The 95% confidence interval for the proportion of significant events, mean shuffled-corrected log odds difference, mean false-positive rates and the adjusted p-value across all methods are available in Figure 6—source data 1.

## Discussion

Previous studies describing the phenomenon of replay have used a wide variety of methods and criteria for replay detection. The most common methods in the last decade have been a rank order correlation (applied to spike trains), and the scoring of decoded trajectories (using a weighted correlation or linear fit) **(Foster, 2017; Tingley and Peyrache, 2020; van der Meer et al., 2020).** However, even within these two classes of methods, there is substantial variation in the parameters and strategies for statistical testing, including the type and number of shuffling procedures, and the use of ripple power or jump distance as a threshold. The diversity in replay methods prevents an unbiased comparison between different replay findings. In particular, while replay detection statistical tests are commonly used with a p-value of 0.05, this value does not necessarily match the actual false-positive detection rate for all methods. This issue is not just limited to when the entire spike train is used to calculate the rank-order correlation (as discussed above), but also applies to all methods using decoded neural activity. The use of either spatially smoothed place fields or spike bursts spanning multiple time bins similarly violates the assumption of independent samples. While the shuffling of neural patterns comprising the replayed trajectory may compensate for non-independent samples, its effectiveness varies substantially, especially given the wide range of parameters used across the published replay literature. Furthermore, while imposing stricter criteria for replay detection would theoretically improve the average fidelity of the events and reduce the false-positive rate, it comes with the caveat of potentially increasing the rejection of true replay events.

Up until now, there has been no unifying approach to quantify and compare the replay detection performance, hindered largely by the phenomenon of replay lacking a ground truth. Here we introduce a novel framework for comparing different detection methods, quantifying the proportion of significant events (quantity), the mean log odds difference (trajectory discriminability), adjusting the threshold used for replay detection according to an empirically-measured false-positive rate of 5%. Using this approach, we make six key observations-**1)** a ripple power threshold is not necessary for RUN replay, but should be applied to POST replay events, **2)** more shuffles, with a preference for pre-decoding shuffles, lead to better replay detection, even after the p-value is adjusted to match the false-positive rate, **3)** without adjusting the p-value, there is a very wide distribution of both the trajectory discriminability and the proportion of significant events detected across methods, **4)** after adjusting the p-value, there is greater similarity observed between methods, however using a weighted correlation replay scoring with three different shuffling procedures or a linear fit replay scoring with two different shuffling procedures still results in the best overall performance for our dataset, **5)** our metric for replay trajectory discriminability yields a similar result across many methods once the p-value is adjusted to fix the false-positive rate at 5%, however the proportion of replay events detected remains more variable, and **6)** regardless of the method used after the p-value is adjusted, preplay events are at chance level for both trajectory discriminability (mean log odds difference of 0) and proportion detected (10% of events across two tracks). Previous studies have used an unadjusted p-value, which in our hands substantially changes the proportion of replay events detected across methods-overly strict methods such as weighted correlation with a jump distance criterion would have a proportion less than 5%, while more lenient methods such as rank order correlation with all spikes included can have a proportion closer to 30%. Therefore, without actually measuring the false-positive rate, one cannot assume that a higher proportion of events relative to the p-value is evidence, on its own, for the existence of replay. In all methods tested, we do not see any evidence of preplay events behaving differently from spurious replay events generated using randomized data. In contrast to this, both POST and RUN replay events have a proportion of significant events substantially above the chance level for detection, for any method used with an adjusted p-value.

Here we have focused on a number of common replay scoring methods and criteria, however there are many variations and nuanced differences between published methods, that were beyond the scope of this study to test. For instance, we did not examine the impact of spatial smoothing (place fields) and temporal smoothing (posterior probability matrix) on replay detection. Based on the desire to maximize the statistical independence of data, we avoided any smoothing in our analysis, as this could increase the false-positive rate of replay detection. Because we did not use spatial or temporal smoothing, we opted to use larger bin sizes for Bayesian decoding (10 cm by 20 ms). It is possible that with smaller bin sizes, replay detection could improve, or yield a different outcome when comparing methodologies. However, as the size of the decoding bin decreases, noise in the decoded trajectory will likely increase or alternatively must be removed with smoothing, in turn potentially increasing the false-positive rate of detection. Replay experiments can often vary along many other parameters (e.g. number of neurons, distribution of place fields, length of track, etc.), and it is possible that these differences could also affect the false-positive rate. As such, it is critical for replay studies to independently verify and report this false-positive rate even for the experiments involving only one spatial context.

It is also important to note that the use of two linear tracks was critical for our current framework, to calculate trajectory discriminability. While this limits the applicability of our approach, with modification, it is possible that this framework might be adapted to single-track data, by calculating trajectory discriminability using a cell-id shuffle to create a virtual second track or by comparing the discriminability across different portions of the track (depending on the experimental design), although this is beyond the scope of our current study.

One key consideration is that due to algorithmic constraints, some sequence detection methods may be biased to detecting certain types of replay trajectories. For instance, a linear fitting method may be more prone to detect linear replay trajectory compared to sigmoidal-shaped or jumpy replay trajectory. On the other hand, a rank-order correlation analysis of spike trains is more relaxed in terms of the rigidity of the temporal structure, and therefore is more likely to include non-linear events with gaps. As such, there may be other factors to consider beyond trajectory discriminability and proportion of detected events when choosing a replay detection method. However, it is interesting to note that despite substantial differences in these algorithms used for replay detection, all methods had a similar trajectory discriminability after adjusting the p-value to a fixed false positive rate of 5%.

Last but not least, in addition to comparing the performance of different replay quantification strategies, this framework can also be applied to compare replay events detected across different behavioral states or experimental conditions. Previously, differences in replay across conditions have been measured using the rate or number of replay events **(Gillespie et al., 2021; Olafsdottir et al., 2015; Xu et al., 2019),** or decoding track bias **(Carey et al., 2019; Tirole & Huelin Gorriz, et al., 2022)**. However, combining these two approaches creates a more comprehensive metric for understanding the difference in replay cross different conditions.

## Supporting information

All supplementary figures

sFigure 4-2 jump distance effect source data 1

Figure 1 source data 1 weighted correlation with two shuffles table

figure 2 source data 1 ripple effect table

figure 3 source data 1 single shuffle table

figure 4 source data 1 shuffle combinations table

figure 5 source data 1 spearmans all spikes versus median spike table

figure 6 source data 1 shuffle-subtracted method comparision table

## ACKOWLEDGMENTS

We thank members of the Bendor Lab for valuable discussion and Aman Saleem for their comments on the manuscript. Rat schematic in Fig. 1A was adapted with permission from SciDraw.io (https://doi.org/10.5281/zenodo.3926277, https://doi.org/10.5281/zenodo.3926237, https://creativecommons.org/licenses/by/4.0). This work was supported by the Biotechnology and Biological Sciences Research Council scholarship (BB/M009513/1) (MTir), the European Research Council starter grant (CHIME) (DB), the Human Frontiers Science Program Young Investigator Award (RGY0067/2016) (DB), and the Biotechnology and Biological Sciences Research Council Research grant (BB/T005475/1) (DB). The Titan Xp used for this research was donated by the NVIDIA Corporation.

## AUTHOR CONTRIBUTIONS

Experiment design: M.Tak, DB, M.H.G, M.Tir

Data collection and processing: M.H.G, M.Tir

Data Analysis: M.Tak

Initial draft: M.Tak, D.B

Manuscript revisions: M.Tak, DB, M.H.G, M.Tir

Both MTir and MHG contributed equally. All authors have given approval to the final version of the manuscript.

## METHODS

All the behavioral and electrophysiological data used in this study were previously described in detail in **Tirole & Huelin-Gorriz, et al. (2022).**

### Behavioral paradigm and electrophysiological recording

Five male Lister-hooded rats (350g-450g) were implanted with 24 independently movable tetrodes, using a custom-made microdrive. In three rats, the microdrive was split into two 12 tetrodes groups, aimed at the right and left dorsal CA1 regions of the hippocampus respectively (AP: −3.48 mm; ML: ±2.4 mm from Bregma). In the remaining 2 rats, right dorsal CA1 was targeted with 16 tetrodes (AP: 3.72 mm, ML: 2.5 mm from Bregma), with the remaining 8 tetrodes targeting visual cortex.

Before the experimental sessions, each animal was trained for approximately two days (30 min sessions) to run back and forth on a linear track. The animal was motivated to run by receiving a reward (chocolate flavoured soy milk) at each end of the track. The training took place in a different room and the track configuration used for training was distinct from the experimental linear tracks.

Each recording session started with a one-hour rest (PRE) epoch in which the rats were placed in a previously habituated rest pot at a remote location. The rest pot was made of circular enclosure (20 cm diameter) surrounded by a 50 cm tall black plastic wall, which prevented the rats from viewing rest of the environment. Following the PRE epoch, the rats went through one of the two protocols described below, for which both were followed by a POST rest epoch in the rest pot:

1. Rats were exposed to 2 novel linear tracks (2 m long).
2. Rats were exposed to 3 novel tracks (2 m long). Data from the first track is not included here to create a two-track experiment consistent with the remaining data.

### Place cell analysis

Place cells were required to have a minimum peak firing rate greater than 1 Hz, based on the unsmoothed ratemap. Spike trains used to compute the place-field were speed filtered (4-50 cm/s).

### Bayesian decoding of spatial location

A naïve Bayesian decoder was used to estimate the brain’s estimation of animal’s position during behavior and virtual position during a replay event, using non-overlapping 20 ms time bins, and 10 cm position bins.

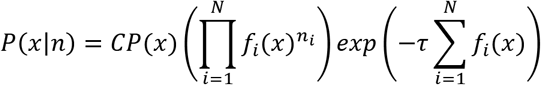

where P(x|n) is the probability of the animal being at a specific position given the observed spiking activity, C is a normalization constant, x is the animal’s position, f_i_(x) is the firing rate of the i^th^ place field at a given location x, and n is the number of spikes in the time window *τ*. The normalization constant was defined as the summed posterior probabilities across both tracks.

### Detection of candidate replay event

Candidate replay events were first selected based on multi-unit activity (MUA), which was smoothed with a Gaussian kernel (sigma = 5 ms) and binned into 1 ms steps. Only events with MUA bursts with a maximum duration of 300 ms and z-scored activity over 3 were selected. Furthermore, ripple power was used as an additional criterion for candidate event selection. Ripple-band (125 - 300Hz) LFP signal was smoothed with a 0.1s moving average filter. Unless it was mentioned otherwise, the candidate events were discarded if the peak ripple power was less than z-score of 3. Moreover, the candidate events were further thresholded such that only events where the animal’s running speed was less than 5 cm/s, at least five active place cells and event duration over 100ms and below 750ms were included for subsequent analyzes. To maximize detection of replay events and avoid discarding minority events due to noisy probability decoding at the beginning or the end of the event, candidate replay events were split into two segments where the midpoint was determined based on the minimum MUA activity in the middle third of the candidate event. Both segments were decoded and analyzed for statistical significance independently with the p-value threshold adjusted to half of the p-value threshold used for ‘full’ event (e.g. p<0.05 → p<0.025). It should be noted that same criteria for candidate event inclusion such as minimum event duration and active place cell number still applied to these ‘half’ events. For this study, RUN replay was also defined as awake replay events when the animals were physically on the linear tracks (with moving speed less than 5cm/s). PRE and POST replay was defined as replay events that took place during rest and sleep periods (not differentiated from each other) within the rest pot before and after experiencing both novel tracks, respectively

### Sequence-based replay scoring

For this study, two different sequence quantification methods were used to quantify the sequenceness of the decoded posterior probability matrix for each event:

#### 1. Weighted correlation

Weighted correlation method calculated the correlation coefficient between the change in decoded posterior probability bins across position and time by weighing each estimated position by its decoded posterior probability:

Weighted mean:

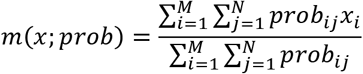

Weighted covariance:

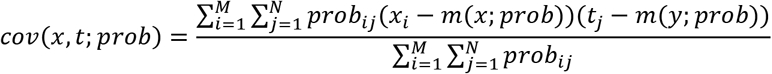

Weighted correlation:

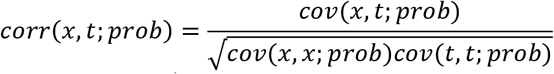

Where *x_i_* is the i^th^ position bin, *t_j_* is the j^th^ time bin and *prob_ij_* is the probability at the position bin *i* and time bin *j*.

#### 2. Linear fitting

The linear fitting method finds the line of best fit that describes the decoded linear trajectory for each candidate event. In practice, this method would calculate all the possible linear kernel at different slopes (from 100 cm/s to 5000 cm/s) and intercept and then sum all the probabilities within 10cm below or above each fitted line as a goodness of fit score:

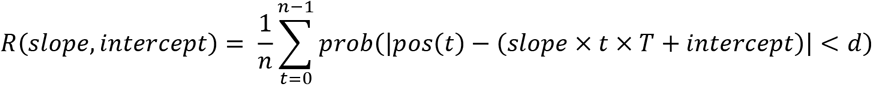

where t is each time bin, prob is the decoded posterior, pos is the position bin, T is the time bin, d the maximum distance from the line of fit.

The combination of slope and intercept that maximizes the posterior decoded probabilities along the potential linear trajectory would be considered as the line of best fit.

Similar sequence-based analyzes were also applied to the cell-id shuffled replay events where the cell identity for each spike train in a candidate replay event was randomized before decoding. This shuffle was designed to preserve the firing statistics across place cells while randomizing the order of place fields along the track, in turn disrupting any relationship with the spike sequences produced during candidate events. For this study, to enhance the accuracy of our false-positive rate calculation, given that false-positive events were generally less frequent than replay events detected during POST and RUN in our original dataset, we generated 3 cell-id shuffled candidate events for each real candidate event (with a new set of random seeds for every event) such that the total number of cell-id shuffled events would equal three times the total number of real candidate events.

Following obtaining a sequence score (either a weighted correlation or linear fit based score) for each event, we then calculated the p-value by comparing the event sequence score relative to the four shuffled distributions, each designed to randomize certain aspects of the sequential place cell firing:

### Pre-Decoding shuffling procedures

**Shuffle 1** : **Spike train circular shuffle,** in which the spike count vectors for each cell were independently circularly shifted in time by a random amount within each replay event, prior to decoding. This was shuffle was designed to degrade the temporal order of the neuronal spiking with minimal disruption of each neuron’s spiking statistics and spatial template.

**Shuffle 2** : **Place field circular shuffle,** in which each ratemap was circularly shifted in space by a random amount of position bins prior to decoding. This shuffle was designed to randomize the preferred firing locations while preserving the temporal structure of the spiking patterns.

### Post-Decoding shuffling procedures

**Shuffle 3** : **Place bin circular shuffle**, in which posterior probability distribution for each time bin (column) was independently circularly shifted by a random amount. This shuffle was designed to shift the decoded position at each time bin while preserving each neuron’s spiking properties.

**Shuffle 4** : **Time bin permutation shuffle**, in which the order of the time bins within each event was permutated randomly. This shuffle was designed to disrupt within-event temporal structure without interfering each neuron’s place field template.

#### 3. Rank-Order Correlation

In addition to a probabilistic decoding based approach, a Rank Order Correlation approach (Spearman correlation coefficient) was also performed directly to the spike train data during candidate replay events using all spikes (method 1) or only the median spike time (method 2) produced by each place cell:

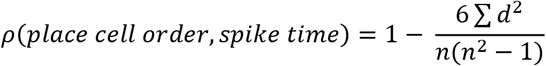

Where d is the difference between the ranks of order of place cells that are active during a given event and the observed spike time, n is the number of spikes included for analysis (i.e. all spikes or only median spike produced by each place cell) and *ρ* is the Spearman’s rank correlation coefficient.

The p-value for Spearman’s *ρ* was computed by comparing the original *ρ* relative to the shuffled distribution where the order of spike was randomly permuted.

### Quantification of significant event proportion and false-positive rate

While standard replay analysis used p<0.05 (i.e. score greater than 95% of the shuffled distribution) as the cut-off point for statistical significance, we calculated the proportion of significant events using p-value thresholds ranging from 0.2 to 0.001. When more than one shuffle type was used together for detection, the sequence score was required to be below the p-value threshold for each shuffle type used, in order for the event to be considered significant. In cases when more than one shuffle was used to calculate the p-value, the highest p-value would be used to represent the sequenceness of each significant replay event. In cases where the candidate replay event was detected as statistically significant for both tracks, (referred as ‘multi-track event’), the candidate event was assigned as both track 1 and track 2 events. However, each multi-track event would be only counted once when calculating the proportion of significant events to avoid double counting. Assuming false-positive rate at 5%, we would expect approximately 10% detection rate (significant events labelled as either track 1 or track 2) with the caveat that this value can potentially decrease in the case where a high number of multi-track events were detected.

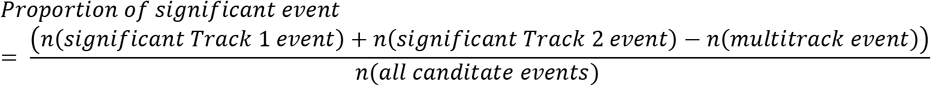

For cell-id shuffled replay events, similar analysis was performed to calculate mean false-positive rate across both tracks.

The mean false-positive rates across both tracks (later used for p-value adjustment) were calculated by adding the number of false-positive events on both tracks divided by 2 (i.e. number of tracks), which was then divided by the total number of candidate events:

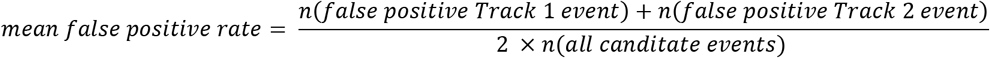

### Sequenceless decoding and quantification of trajectory discriminability

In order to cross-validate the quality of replay events detected based on sequence-based approach, we used an independent metric that measure reactivation bias based on sequenceless decoding **(Carey et al., 2019; Tirole & Huelin Gorriz, et al., 2022).** Only cells with stable place fields in the first and second half of the behavioral episode on both tracks (peak in-field firing rate >1 Hz) were included in our sequenceless decoding analysis. Prior to Bayesian decoding, ratemaps for Track 1 and Track 2 were concatenated as a single matrix [*nCell* × (*nPos*(*T*1) + *nPos*(*T*2))], where nPos is the number of position bin on each track (i.e. 20 × 10cm position bins for both tracks). For a given replay event, the decoded posterior probabilities for each time bin were normalized across all position bins from both tracks to sum to 1. We quantified the logarithmic ratio between the two tracks, and then z-scored this ratio relative to a shuffled distribution computed by Track ID shuffle. For each place cell in the track ID shuffle, its place field templates for track 1 and track 2 were randomly assigned (i.e. each cell’s ratemap was either swapped or not swapped). Then, we quantify the trajectory discriminability in terms of the difference in mean log odds between track 1 and track 2 replay events, originally detected using sequence-based replay detection methods, to cross-validate the quality of replay content.

### p-value adjustment based on mean false-positive rate across both tracks

After calculating the mean false-positive rate across both tracks at p-value threshold ranging from 0.2 to 0.001, we then identify the p-value that leads to the mean false-positive rate closest to 0.05 for a given method. The log odds difference and proportion of significant events detected at the adjusted p-value threshold would be used for the subsequent method comparisons.

### Statistics

#### Bootstrapping for method comparisons

To determine if the differences between two methods (i.e. mean log odds or proportion of significant events) were statistically significant, we used a bootstrapping procedure where candidate replay events were resampled with replacement 1000 times. This created bootstrapped distributions of log odds differences, the proportion of significant events detected and false-positive rates at each p-value threshold ranging from 0.2 to 0.001.

When comparing the proportion of significant events and log odds difference at the original p-value < 0.05, we calculated the 95% confidence interval for both metrics obtained at a p-value < 0.05. When comparing the proportion of significant events and log odds difference at the adjusted p-value, we calculated the 95% confidence interval for both metrics obtained at the p-value when the mean false-positive rate was closest to 5%. The difference between the two bootstrapped distributions was only considered statistically significant when the 95% confidence intervals did not overlap.

#### Bootstrapping for log odds significance

To determine if the log odds difference of the replay events detected during PRE, RUN and POST by a given method was statistically significant from chance level trajectory discriminability, we calculated the confidence interval for the difference between bootstrapped distribution of the original replay data and the cell-id shuffled replay data. The mean difference between two bootstrapped distributions were only considered statistically significant when the 95% confidence interval did not overlap with 0.

