## Supplementary material for "Evaluating hippocampal replay without a ground truth": All supplementary figures

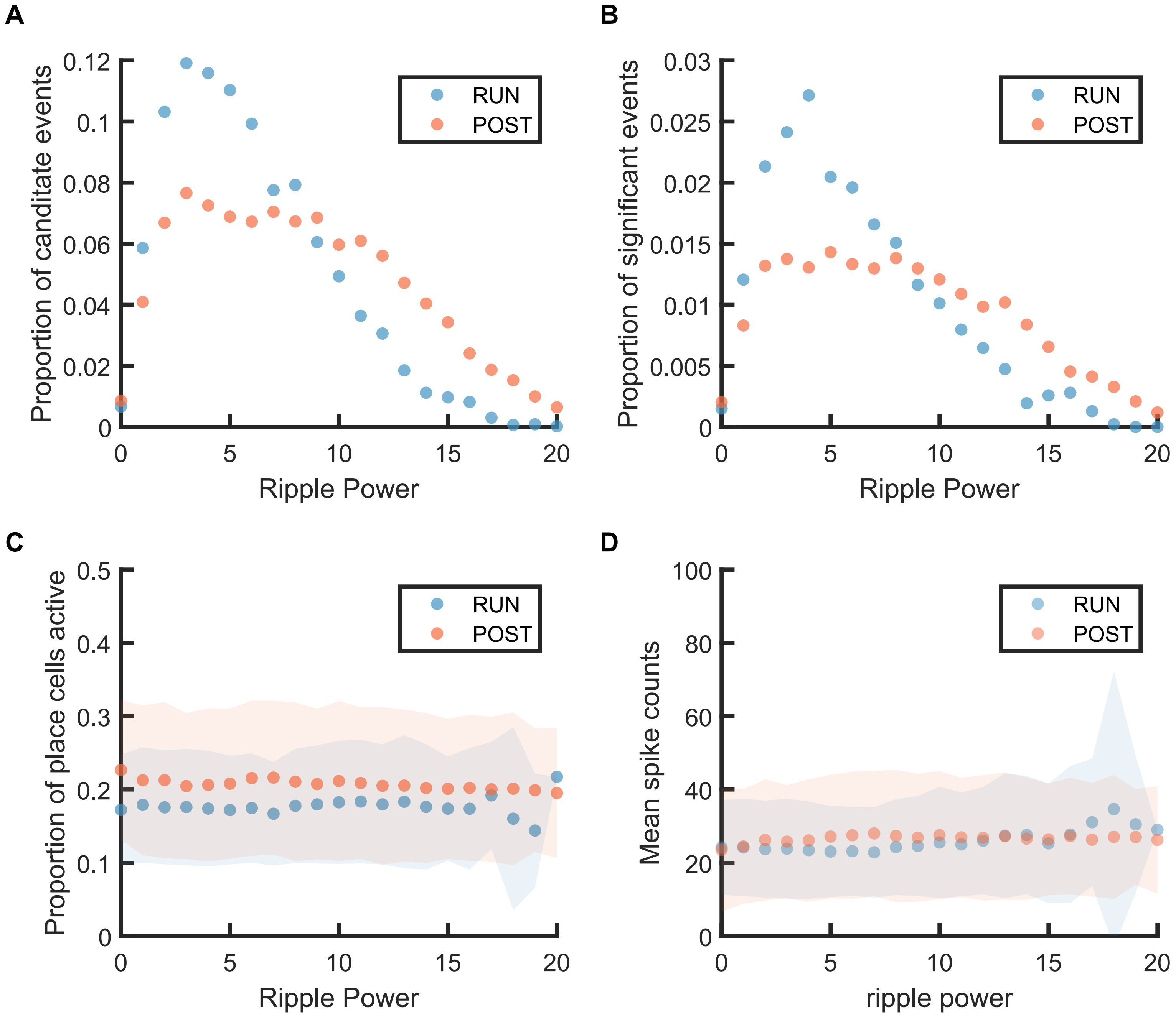


**Figure S2-1 The replay event distribution, proportion of place cells active and total spikes during replay event at different ripple powers. (A)** The distribution of candidate replay events across different ripple powers **(B)** the proportion of significant events (out of all candidate events) at p value ≤ 0.05 at different ripple powers. **(C)** The mean proportion of place cells active during significant replay events. The shaded region indicated the standard deviation of the number of active place cells at each ripple power range. **(D)** The mean number of spikes fired by the active place cells during significant replay events. The shaded region indicated the standard deviation of the spike count at each ripple power range.

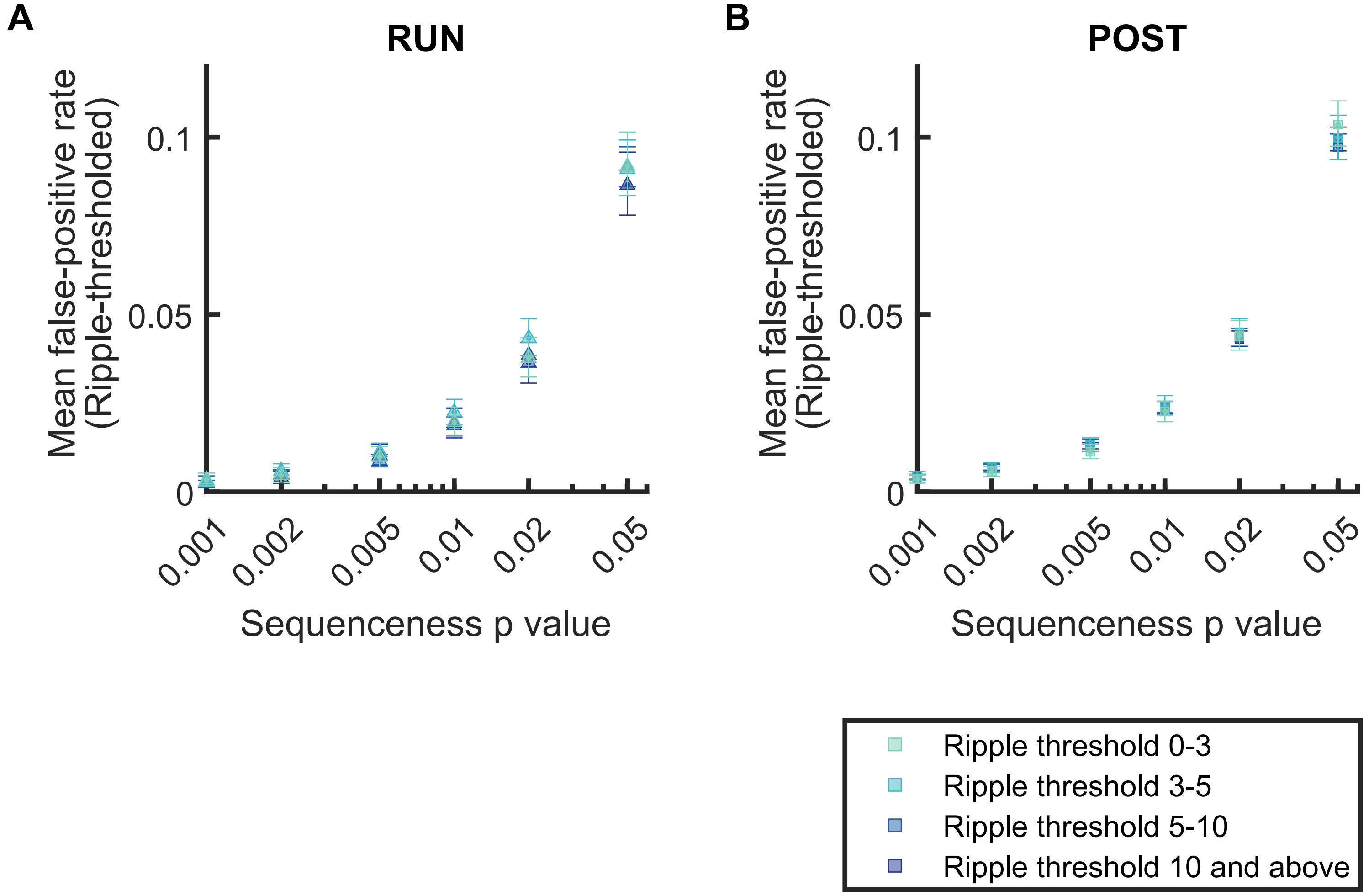


**Figure S2-2 The mean false-positive rate across both tracks for replay events detected with different ripple power range. (A,B)** The mean false-positive rate calculated at different P-value thresholds (i.e. 0.05, 0.02, 0.01, 0.005, 0.002, 0.001) as ripple power increased (i.e. 0-3, 3-5, 5-10,10 and above). The error bar indicated the 95% bootstrap confidence interval **(A)** Replay events detected during RUN **(B)** Replay events detected during POST

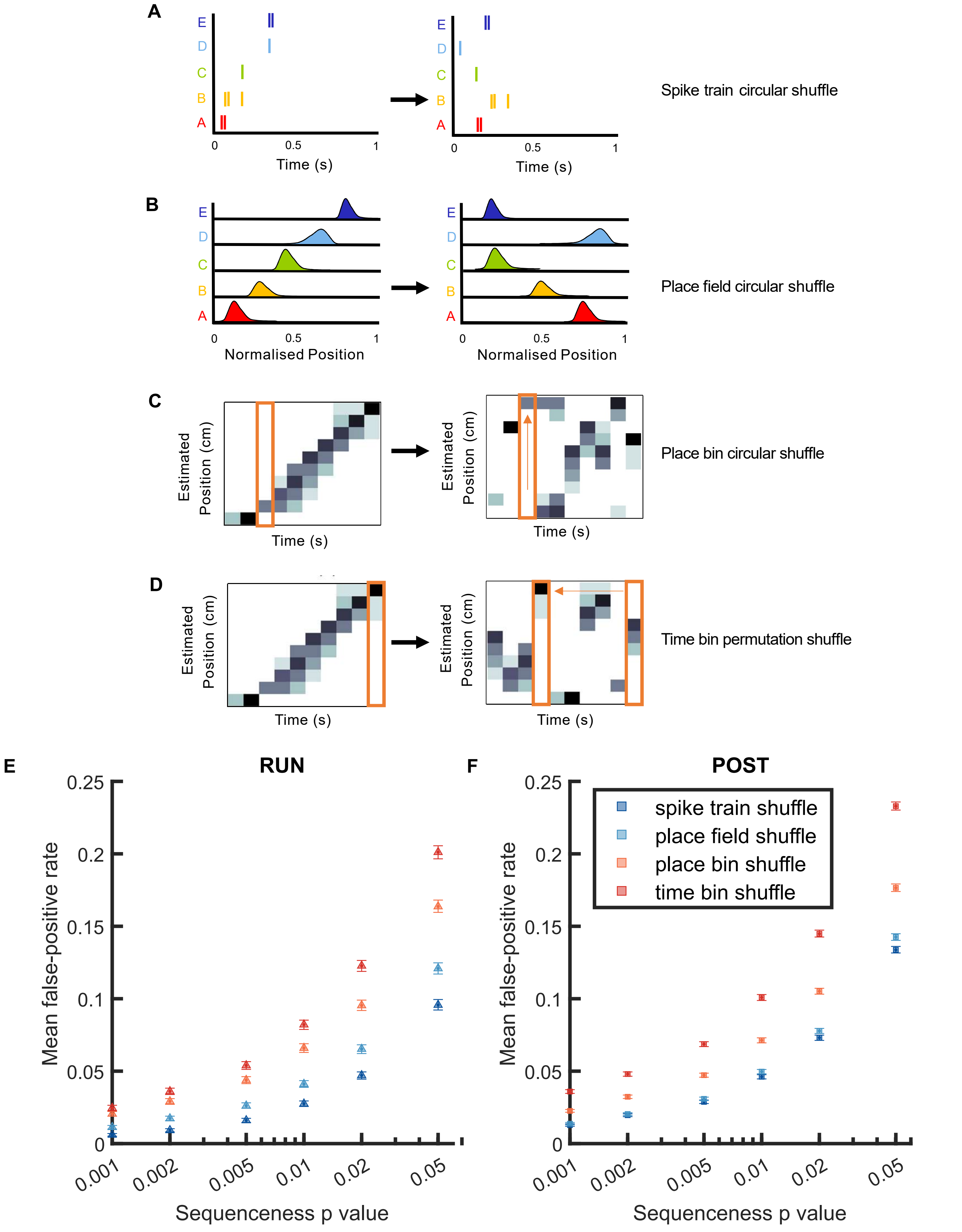


**Figure S3-1 The mean false-positive rate across both tracks for replay events detected using different shuffling methods. (A)** Spike train circular shuffle (dark blue) in which each cell's spike train was independently circularly shifted in time by random amount within each replay event. **(B)** Place field circular shuffle (light blue) in which each cell's ratemap was circularly shifted in space by a random amount. **(C)** Place bin circular shuffle (orange) in which the posterior probability distribution for each time bin was independently circularly shifted by a randomly amount. **(D)** Time bin permutation shuffle (red) in which the order of the time bin was permuted randomly. **(E,F)** The mean false positive rate calculated at different p value thresholds (i.e. 0.05, 0.02, 0.01, 0.005, 0.002, 0.001) using four shuffles. The error bar indicated the 95% bootstrap confidence interval **(E)** Replay events detected during RUN **(F)** Replay events detected during POST

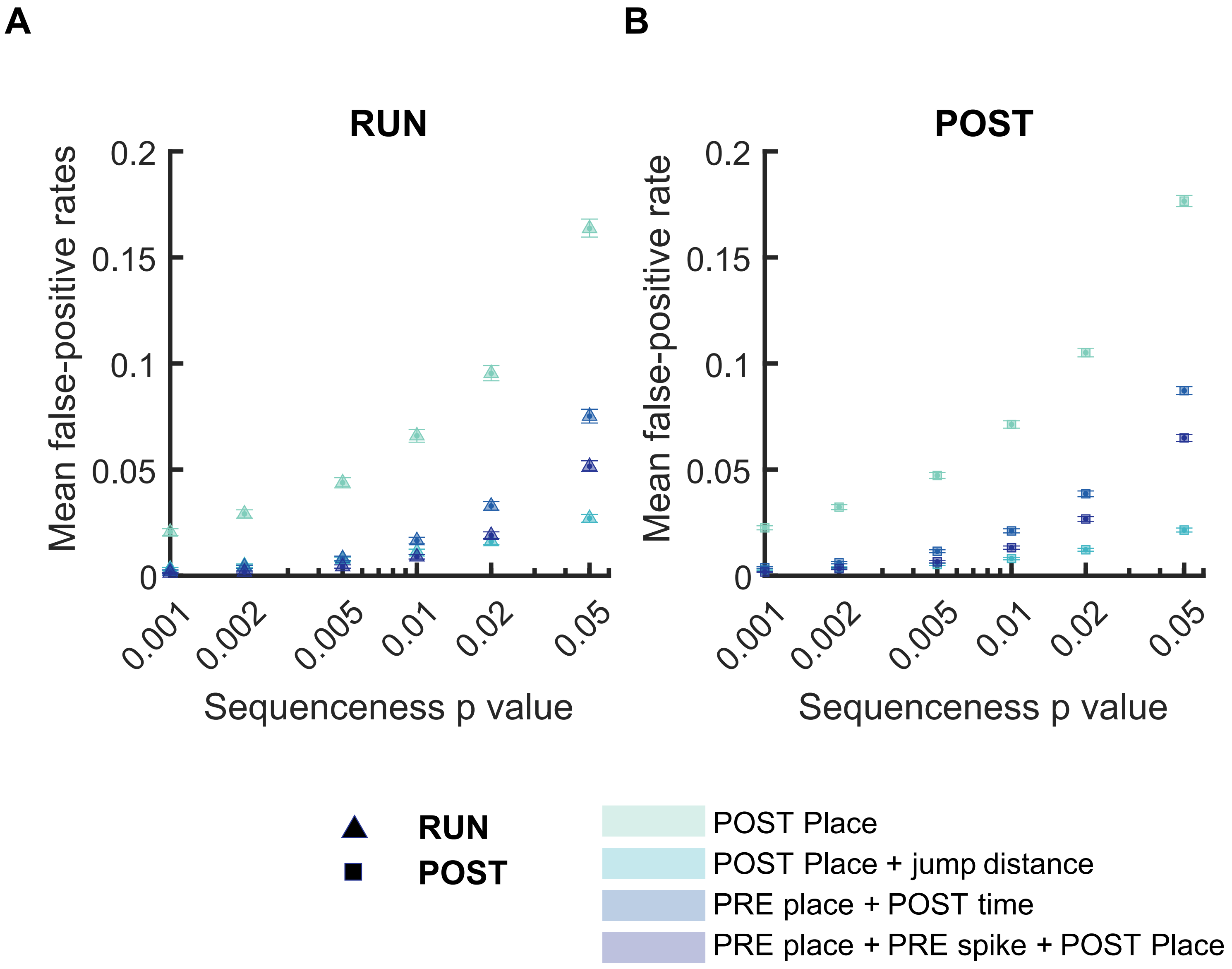


**Figure S4-1 The mean false positive rate across both tracks for replay events detected using different detection criteria. (A,B)** The mean false positive rate calculated at different p value thresholds (i.e. 0.05, 0.02, 0.01, 0.005, 0.002, 0.001) using four different detection ceriteria: **(1)** Only a place bin circular shuffle **(2)** a place bin circular shuffle with jump distance threshold (40% of the track length) **(3)** a place field circular shuffle and a time bin permutation shuffle, **(4)** a place bin circular shuffle, a spike train circular shuffle and a place bin circular shuffle. The error bar indicated 95% bootstrap confidence interval. **(A)** Replay events detected during RUN **(B)** Replay events detected during POST.

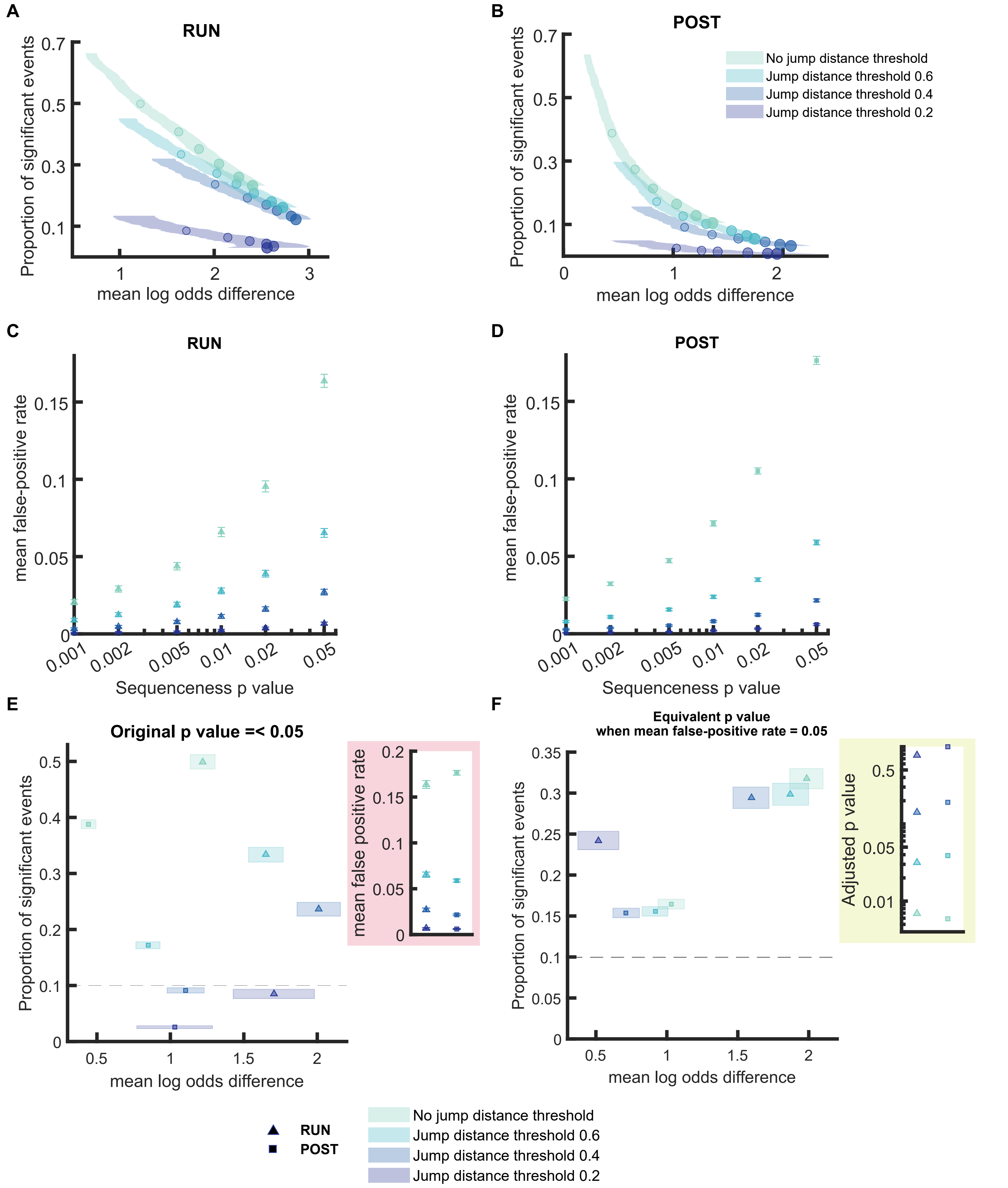


**Figure S4-2 Comparison of the replay detection performance when applying different jump distance thresholds (A,B)** The proportion of significant events and mean log odds difference at different p value thresholds (0.2 to 0.001) using place bin circular shuffle plus jump distance threshold ranging from 0.2 to 1 (normalised track length). The shaded region indicated 95% bootstrap confidence interval. The six dots with increasing colour intensity for each distribution represented the data at p value threshold 0.05, 0.02, 0.01, 0.005, 0.002 and 0.001. **(A)** Replay events detected during RUN **(B)** Replay events detected during POST. **(C,D)** The mean false positive rate calculated at different p value thresholds (i.e. 0.05, 0.02, 0.01, 0.005, 0.002, 0.001) using four different jump distance threshold. The error bar indicated 95% bootstrap confidence interval. For jump distance, due to the extremely low false positive even at p value < 0.2, we quantified the replay detection performance up until p value < 1. **(C)** Replay events detected during RUN **(D)** Replay events detected during POST. **(E,F)** The proportion of significant events and mean log odds difference at **(E)** unadjusted p value ≤ 0.05 and **(F)** adjusted p value when mean false positive rate = 0.05. The shaded box indicated 95% bootstrap confidence interval. The triangle symbol was used to represent replay events during RUN and the square symbol was used to represent replay events during POST. The dashed line represented the approximate chance level at mean false-positive rate of 5%

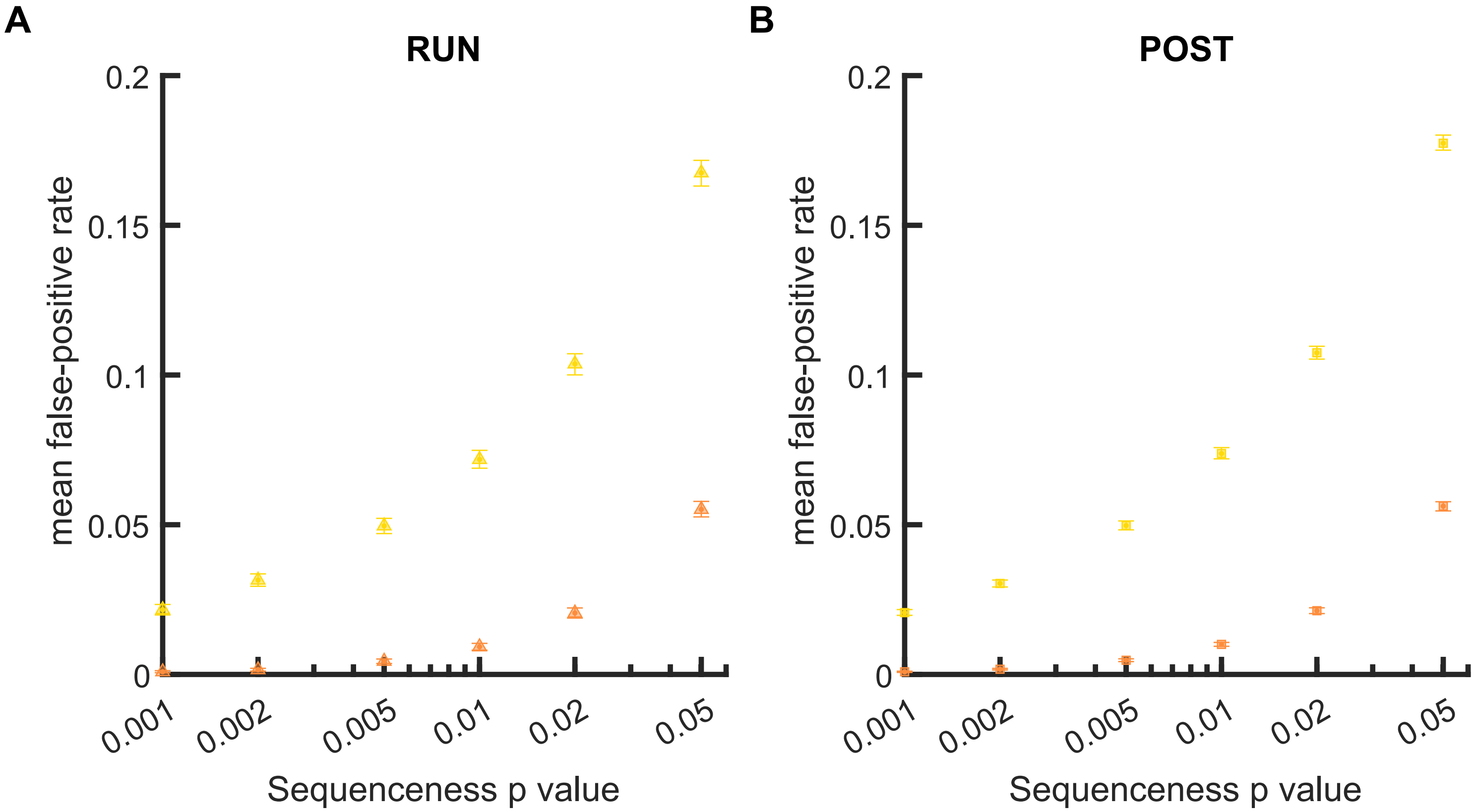


**Figure S5-1 The mean false positive rate across both tracks when all spikes or median spike fired by each place cell was included for Spearman's rank-order based analysis. (A,B)** The mean false positive rate calculated at different p value thresholds (i.e. 0.05, 0.02, 0.01, 0.005, 0.002, 0.001) when **(1)** all spikes or **(2)** only the median spike fired by each place cell was included for rank-order based replay analysis. The error bar indicated the 95% bootstrap confidence interval. **(A)** Replay events detected during RUN **(B)** Replay events detected during POST.

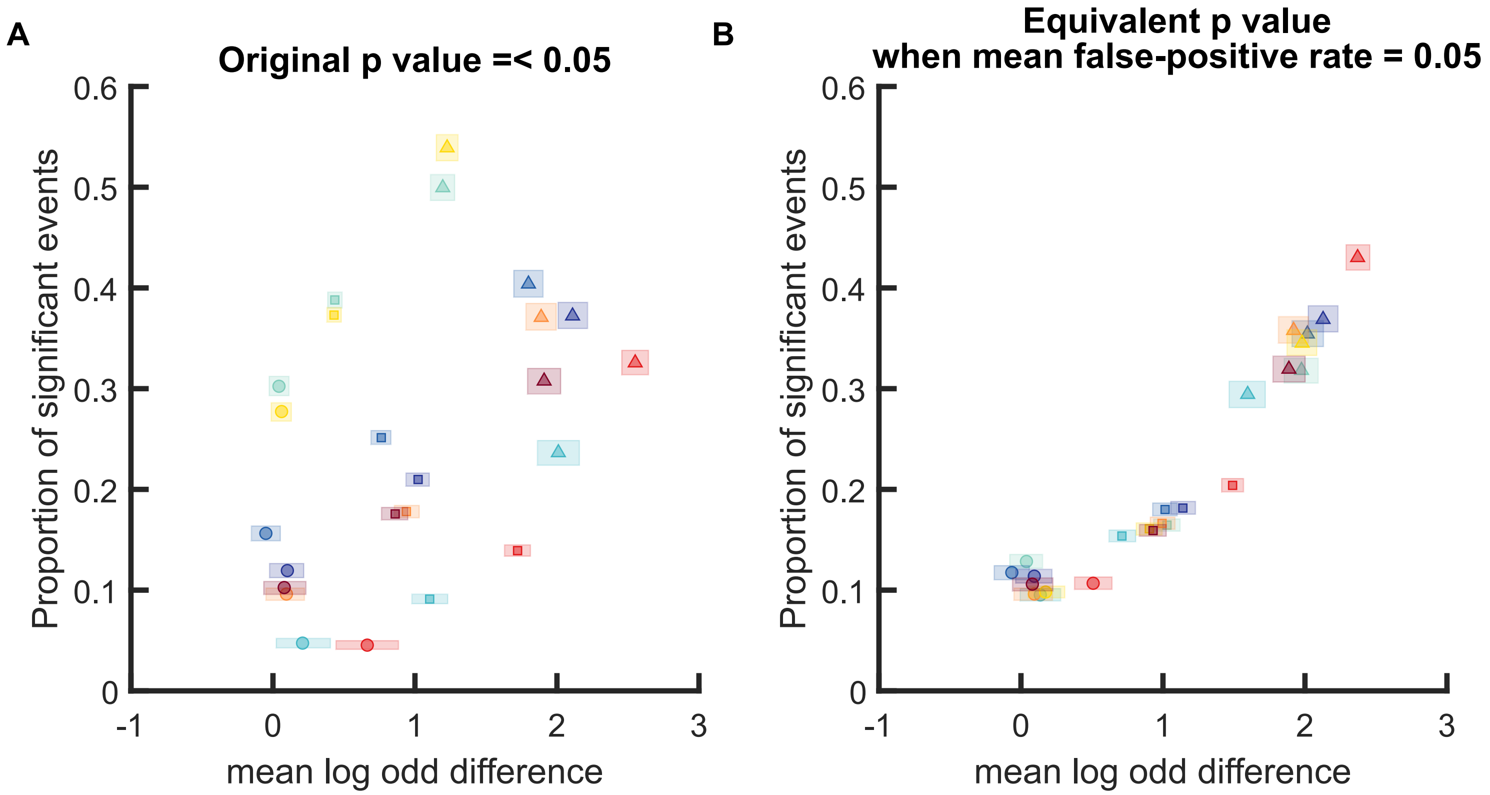


**Figure S6-1. Comparison of different replay detection methods for replay events during PRE, RUN and POST when log odds difference was not shuffle-subtracted (A,B)** The proportion of significant events and original mean log odds difference at different p value thresholds (0.2 to 0.001) using a range of different methods: **(1)** Weighted correlation with place bin circular shuffle **(2)** Weighted correlation with place bin circular shuffle and jump distance threshold at normalised track length 0.4 **(3)** Weighted correlation with place field circular shuffle and time bin permutation shuffle, **(4)** Weighted correlation with place field circular shuffle, spike train circular shuffle and time bin permutation. **(5)** Spearman rank-order based correlation using only median spike fired by each place cell **(6)** Spearman rank-order based correlation using all spikes fired by each place cell **(7)** Liner fitting with place bin circular shuffle **(8)** Liner fitting with place field circular shuffle and time bin permutation shuffle. The shaded region indicated 95% bootstrap confidence interval. The six dots with increasing colour intensity for each distribution represented the data at p value threshold 0.05, 0.02, 0.01, 0.005, 0.002 and 0.001. **(A)** Replay events detected during RUN. **(B)** Replay events detected during POST. **(C,D)** The proportion of significant events and mean log odds difference at **(C)** unadjusted p value $\leq 0.05$ and **(D)** adjusted p value when mean false positive rate = 0.05. The shaded box indicated 95% bootstrap confidence interval. The circle symbol was used to represent replay events during PRE. The triangle symbol was used to represent replay events during RUN and the square symbol was used to represent replay events during POST.

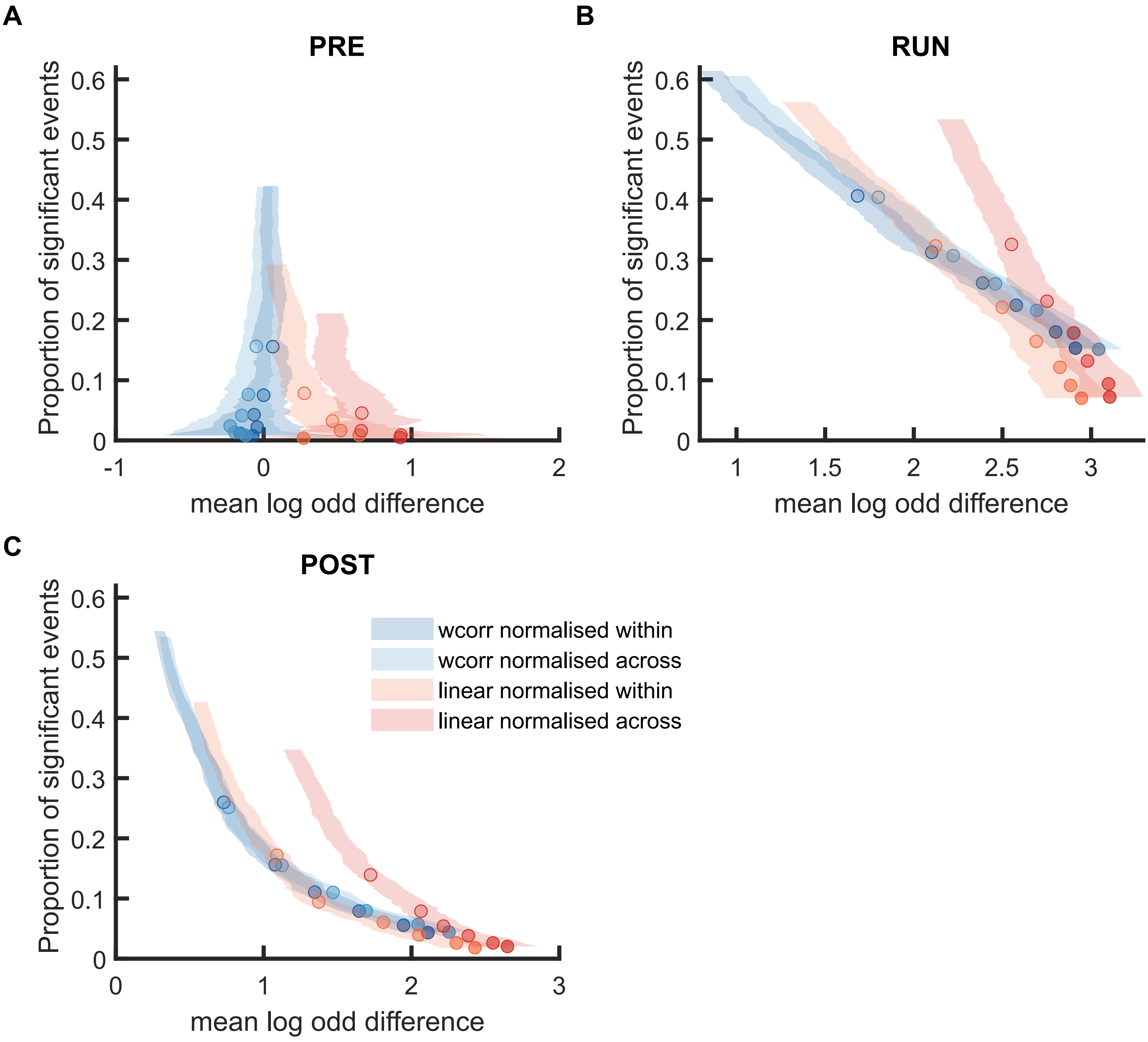


**Figure S6-2 Comparison of mean log odds difference using weighted correlation and linear fitting when the posterior probabilities were normalised within track or cross track (A,B)** The proportion of significant events and mean log odds difference at different p value thresholds (0.2 to 0.001) when using weighted correlation and linear fitting where the posterior probabilities were normalised within track or cross track. The shaded region indicated 95% bootstrap confidence interval. The six dots with increasing colour intensity for each distribution represented the data at p value threshold 0.05, 0.02, 0.01, 0.005, 0.002 and 0.001. **(A)** Replay events detected during RUN **(B)** Replay events detected during POST.

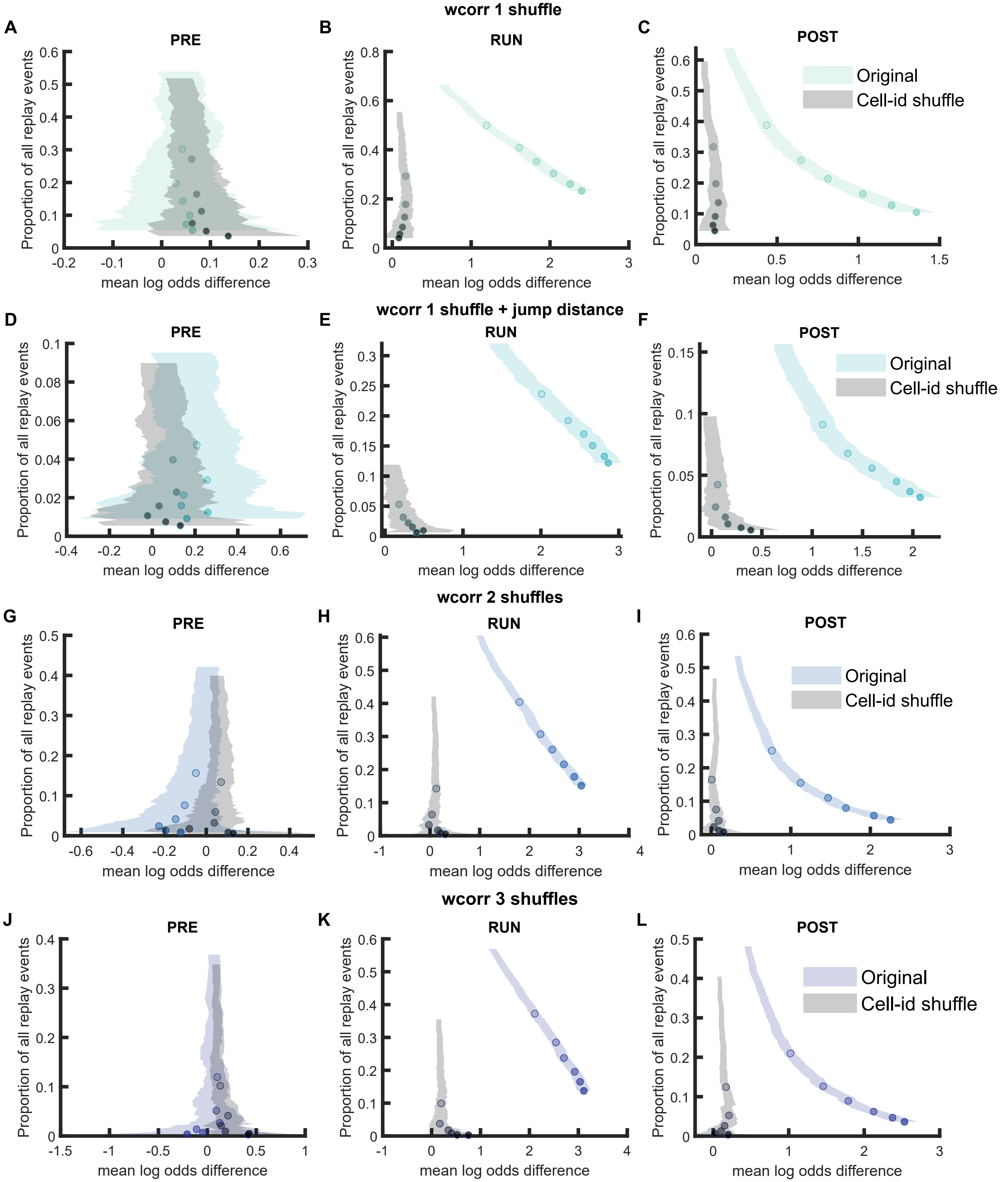


**Figure S6-3 Comparison of replay detection performance using weighted correlation for replay events during PRE, RUN and POST (A-J)** The proportion of significant events and mean log odds difference at different p value thresholds (0.2 to 0.001) when using weighted correlation with different detection criteria. **(A-C)** Only post-decoding place bin circular shuffle. **(D-F)** post-decoding place bin circular shuffle with jump distance threshold at normalised track length 0.4 **(G-I)** Pre-decoding place field circular shuffle and Post-decoding time bin permutation shuffle, **(J-L)** Pre-decoding place field circular shuffle, Pre-decoding spike train circular shuffle and Post-decoding time bin permutation. The shaded region indicated 95% bootstrap confidence interval. The six dots with increasing colour intensity for each distribution represented the data at p value threshold 0.05, 0.02, 0.01, 0.005, 0.002 and 0.001. **(A,D,G,J)** Replay events detected during PRE. **(B,E,H,L)** Replay events detected during RUN. **(C,F,I,L)** Replay events detected during POST

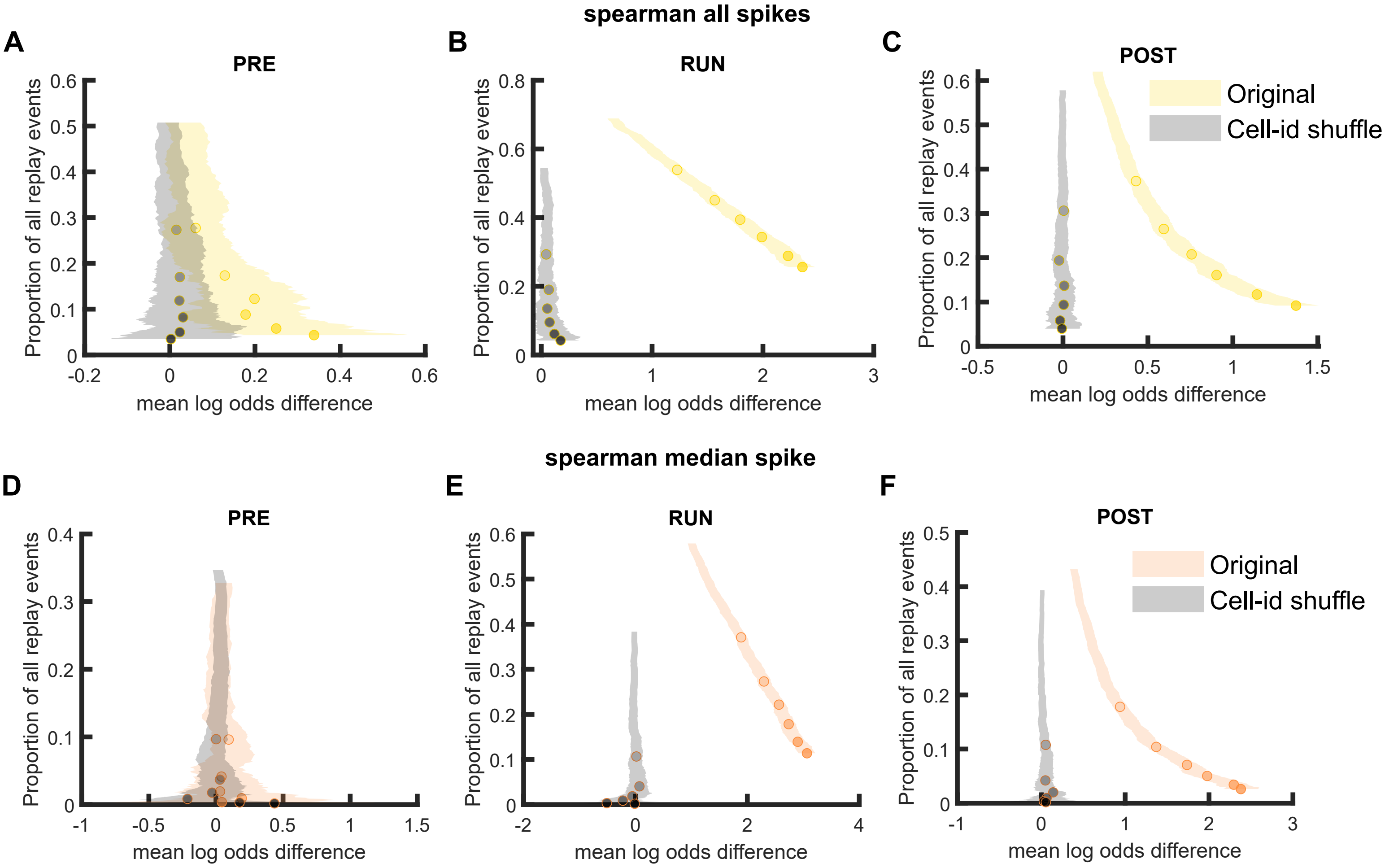


**Figure S6-4 Comparison of replay detection performance using Spearman's rank-order based correlation for replay events during PRE, RUN and POST (A-F)** The proportion of significant events and mean log odds difference at different p value thresholds (0.2 to 0.001) when **(A-C)** all spikes or **(D-F)** only the median spike fired by each place cell was included for rank-order based replay analysis. The shaded region indicated 95% bootstrap confidence interval. The six dots with increasing color intensity for each distribution represented the data at p value threshold 0.05, 0.02, 0.01, 0.005, 0.002 and 0.001. **(A,D)** Replay events detected during PRE. **(B,E)** Replay events detected during RUN. **(C,F)** Replay events detected during POST.

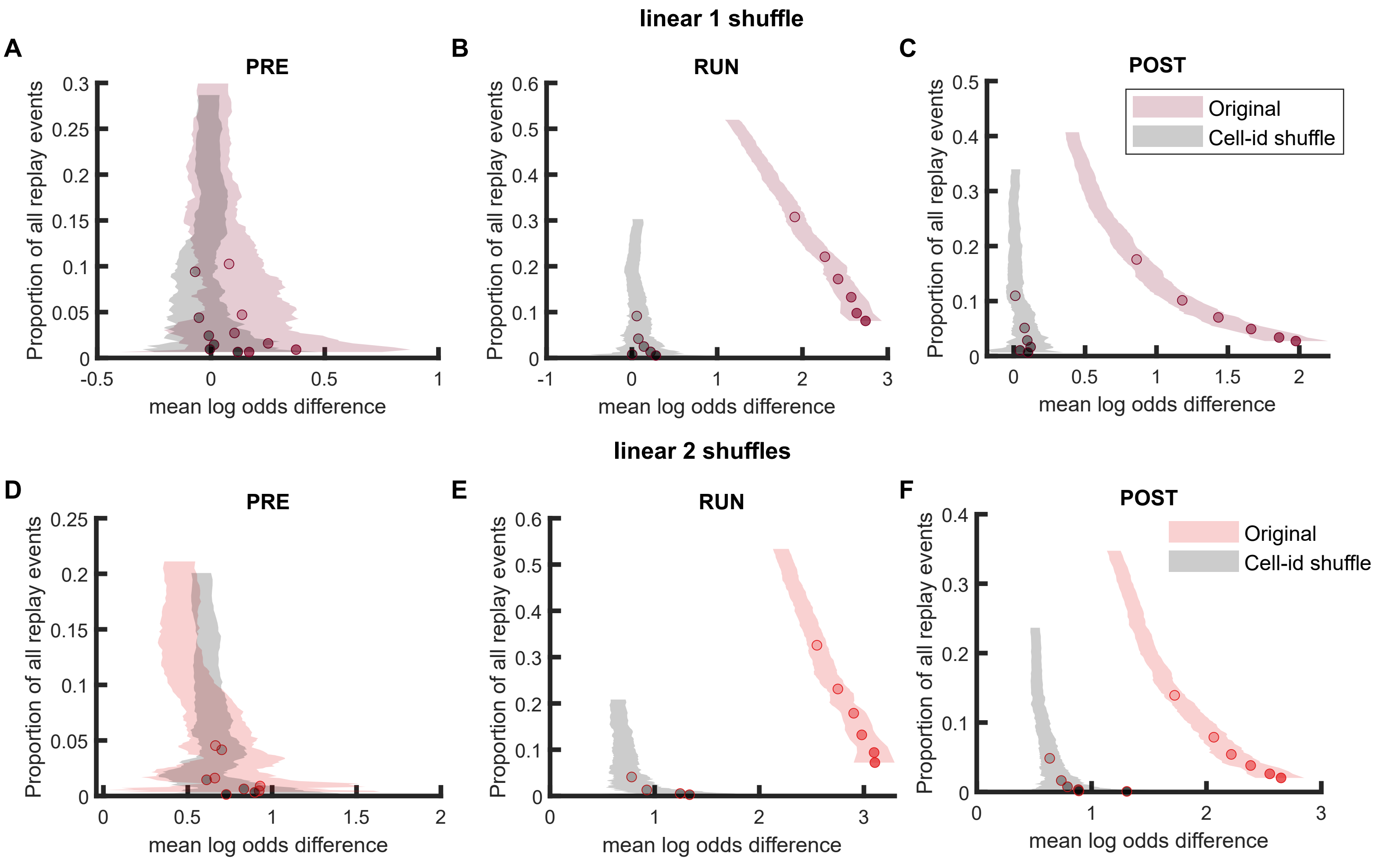


**Figure S6-5 Comparison of replay detection performance using linear fitting for replay events during PRE, RUN and POST (A-F)** The proportion of significant events and mean log odds difference at different p value thresholds (0.2 to 0.001) when using linear fitting approach with **(A-C)** only post-decoding place bin circular shuffle or **(D-F)** both pre-decoding place field circular shuffle and Post-decoding time bin permutation shuffle. The shaded region indicated 95% bootstrap confidence interval. The six dots with increasing colour intensity for each distribution represented the data at p value threshold 0.05, 0.02, 0.01, 0.005, 0.002 and 0.001. **(A,D)** Replay events detected during PRE. **(B,E)** Replay events detected during RUN. **(C,F)** Replay events detected during POST
